# Single-cell analysis of microglial transcriptomic diversity in subarachnoid hemorrhage

**DOI:** 10.1101/2021.12.27.474223

**Authors:** Junfan Chen, Lei Sun, Hao Lyu, Zhiyuan Zheng, Huasheng Lai, Yang Wang, Yujie Luo, Gang Lu, Wai Yee Chan, Yisen Zhang, Xinyi Chen, Zhongqi Li, Ho Ko, Kwok Chu George Wong

## Abstract

**Background:** Subarachnoid hemorrhage (SAH) is a severe stroke and the advanced treatment for SAH is still limited. Recent studies have shown that microglia-mediated neuroinflammation plays a critical role in the pathogenesis of SAH. Microglia can transform their states in response to central nervous system injury. However, the transcriptomic features of microglia remained unknown in SAH. Recent developed single-cell RNA sequencing (scRNA-seq) provides a possible way to solve this problem.

**Methods:** Endovascular perforation (EVP) murine SAH model was established to reproduce experimental SAH. Microglia states are examined with immune staining and quantitate analysis. Post-SAH microglial single-cell suspension were harvest and sequenced using 10X scRNA-seq platform. Then, the detailed single-cell transcriptomic characterization of post-SAH microglia were analyzed with bioinformatics.

**Results:** Transcriptional analysis revealed at least ten diverse microglial subgroups, including SAH-associated microglia (SAM), inflammatory-associated microglia (IAM) and proliferation-associated microglia (PAM), which all exhibit distinct marker gene expression patterns. Microglia subsets interaction reveals the functional relationship between elevated signaling pathways and microglial sub-populations in SAH. Receptor-ligand pair analysis revealed that complex inter-cellular interactions exist between the microglia subsets and other cell types, and indicated that microglia are important mediators of neuroinflammation after SAH. Integrated analysis with normal microglia further proved the existence of these microglia subpopulations and different gene markers associated with SAH were clarified.

**Conclusions:** Collectively, we first report the single-cell transcriptome of post-SAH microglia and found specific biomarkers related to the neuroinflammation in SAH. These results enhanced our understanding of the pathological mechanisms of microglial response to SAH, and may guide future development of SAH monitoring methods and therapeutics.

## 1. Background

Subarachnoid hemorrhage (SAH) is a type of stroke with high morbidity and mortality ^1–3^. Patients who survived from SAH may suffer from severe complications, such as motor disability and cognitive impairment, which lead to serious decline in quality of life ^4–6^. Recently, increasing evidence points to a pivotal role of neuroinflammation in the pathogenesis of SAH ^7–10^. Microglia are brain parenchymal macrophages, the activation of which mediates neuroinflammation responses to pathological states in aging, neurodegenerative diseases, and cerebrovascular accidents ^4, 11–13^. However, the molecular characteristics of microglia activation in SAH-induced neuroinflammation remain unclear.

Activated microglia are characterized by morphological and functional changes from homeostatic to reactive states, presumably driven by the associated complex expression changes ^11^. Depending on the disease context, activated microglia may proliferate and accumulate at sites of pathology, exhibit characteristics of enhanced phagocytosis of cellular debris, and/or produce proinflammatory cytokines and chemokines ^14^. Simulating the aneurysm rupture and subsequent cascades, the endovascular perforation (EVP) animal model is considered an ideal mimic for both the hemorrhagic process and the subsequent pathophysiological changes in SAH ^15–18^. In the murine EVP model, activation and polarization of microglia have been observed ^19–22^. Uncovering the transcriptomic features of microglia would not only help us determine how microglia assume their reactive states in SAH, but also guide us on how microglia subgroups may interact with each other, and other brain cell types, to serve protective functions or exacerbate brain injury.

In this work, we performed transcriptomic profiling of microglia in the EVP mouse model of SAH at the single-cell level. We report the identification of diverse microglial transcriptomic states and subsets biomarkers, that can be classified into SAH-associated microglia (SAM), inflammation-associated microglia (IAM), and proliferation-associated microglia (PAM) et al. The transcriptional features of these microglia subtypes in SAH are each characterized by the upregulated expression of unique panels of genes, and distinct from previously reported disease-associated microglia subtypes. Functional pathway enrichment and cell-cell interaction analyses further substantiate the importance of microglia in driving post-hemorrhage neuroinflammatory responses. Integrated analysis of SAH-associated and normal microglia transcriptomics further support the transcriptional differences. Biomarkers in SAH microglia subpopulations are considered to be closely related to the microglial activation, pathological process and prognosis of SAH, which will be important targets for future research. Collectively, these results provide new insights into the transcriptomic and functional diversity of microglia in SAH, and the mechanisms of associated neuroinflammation.

## 2. Methods

All the detailed methods can be found in **Supporting Information**.

## 3. Results

### 3.1. Microglia become activated and accumulate in the early state of SAH

We first established a mouse EVP model of SAH. Overall, the mortality of our SAH model was 19.4%, in line with previous reports of murine SAH models [8] and similar to the observed human SAH in-hospital mortality ^4, 23^. On the third day after the EVP procedure, the SAH group had the greatest weight loss and motor ability change on the holding time test, while the normal and sham control groups did not exhibit changes in these measures (**Figures S1-A**). Throughout the study, animal subjects with moderate severity based on the modified Bederson score and motor score were selected for further experimentation (**Figure S1-A**, also see **Methods**). With immunohistochemistry, we noted prominent microglia accumulation in the cortex adjacent to the perforated site (CAPS) areas (**Figure 1-B**, **Figure S1-B, S1-C and S1-D**). The accumulated microglia exhibited more amoeboid-shaped morphology, consistent with activated states (**Figure S2, S3 and S4-A**).

**Figure legend 1:**
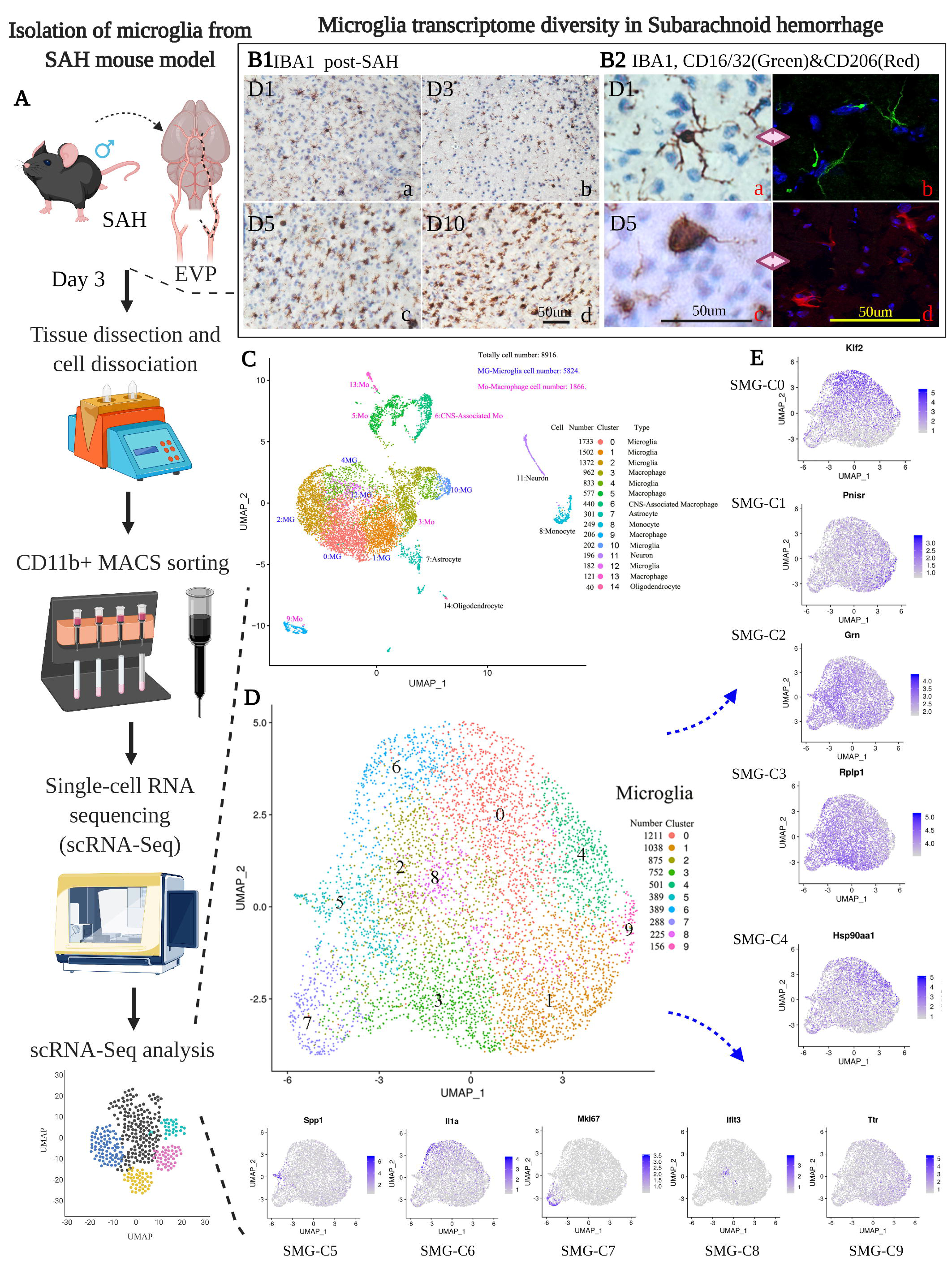
A: Study design of single-cell microglial transcriptomic in SAH; B1-Microglia immunohistochemistry (IHC) staining in D1, D3, D5 and D10 at cortex adjacent to perforate site (CAPS); B2-IHC staining of Ramified and activated microglia and immunofluorescence staining of CD16/32 (Green) and CD206 (Red) of microglia. C: UMAP results of isolated cells; D: UMAP of SAH microglia (SMG); E: Biomarkers of 10 clusters in SMG.

Based on these results, we proceeded to perform single-cell transcriptomic profiling of microglia and other brain cell types on post-SAH day 3 (**Figure 1A**), to investigate the genome-wide transcriptional changes in the early brain injury stage (EBI) of SAH. To increase the throughput of microglia isolation, we carried out magnetic-activated cell sorting (MACS) of CD11b+ cells with high purity (81.5%) and viability (97%) for single-cell RNA sequencing (**Figures S4-B**). We obtained a total of 13,194 single-cell transcriptomes from 4 SAH mouse brains, among which 8,916 (67.6%) were retained for further analysis (**Excel S1**) after quality-control filtering (**Excel S1, Figures S4-C**, see **Methods**).

### 3.2. Single-cell RNA sequencing revealed diverse microglial states after SAH

Based on cell type-specific marker gene expression patterns, we classified the single-cell transcriptomes into microglia (*n* = 5,824), central nervous system (CNS)-associated macrophages (*n* = 440), macrophages (*n* = 1,866), astrocytes (*n* = 301), monocytes (*n* = 249), neurons (*n* = 196) and oligodendrocytes (*n* = 40) (**Figure 1-C and Figure S5**). Upon visualization with dimensionality reduction by Uniform Manifold Approximation and Projection (UMAP), the microglia transcriptomes did not fall into distinct clusters but rather spanned a continuum (**Figures 1-C**). Nonetheless, we employed unsupervised clustering to classify the microglia into ten distinct clusters (**Figure 1-D**), as this permitted us to examine the transcriptomic features of their different states and investigate the molecular diversity of microglia post-SAH. These clusters constituted 2.7 – 20.8% of all microglia (**Figure 1-D**), and each were characterized by a panel of highly expressed genes (**Figure 1-E**, **Figure 4** and **Figure S8, S9 and S10, Supporting information**). We termed these microglia clusters obtained from SAH models SAH microglia (SMG) clusters. We then examined the molecular characteristics of the SMG clusters in detail, to infer their potential functional specializations (see violin plots and dot plots in **Figure S8**, **Figure S9, and Figure S10** for the top 10 expressed genes in each of the 10 SMG clusters, also see **Figure 4** and **Excel S2**)

#### 3.2.1. Homeostatic microglia: SMG-C0, 1, 3 and 4

Among the SMG clusters, several SMG clusters appeared to serve important homeostatic functions (SMG-C0, 1, 3 and 4). SMG cluster 0 (SMG-C0) was characterized by a high expression of immediate early genes encoding transcription factors (*Jun, Junb, Jund, Fos, Egr1, ler5, Klf6, Klf2 and Atf3 etc.*) ^24^. By pathway enrichment analysis, the expression of SMG-C0 was enriched with the negative regulation of the cellular metabolic process, regulation of the apoptotic process, cellular response to stress and response to organic substances (**Figure S6-A, Figure S9-A**). SMG-C1 represented a microglial cluster that expresses high levels of homeostatic genes, including *P2ry12, P2ry13, Siglech, Gpr34, Selplg* and *Pmepa1*. SMG-C1 also has a high expression of *Clec4a3*, which is usually expressed in resident tissue macrophages ^25^. (**Figure S7-B, Figure S9-B**) In the SMG-C3 subpopulation, high expressions of ribosomal were found, including *Rplp1*, *Rplp0*, *Rpl41*, *Rps27a* and *Rps23 etc.* (**Figure S7-A, Figure S10-A**).

SMG-C4 has high transcript counts for genes encoding important transcription factors related to microglia reactive changes (e.g., *Mafb, Btg2* and *Txnip*). They also have high expression of microglial homeostasis genes (*P2ry12, Selplg* and *P2ry13*), HSP70 family and *Malat1*. *Mafb* has been recognized as an important transcription factor especially in adult microglia. *Mafb* is crucial for maintaining a healthy homeostatic microglial phenotype by inhibiting inflammatory changes and proliferation ^26^. Microglia-specific knockout of *Mafb* could lead to disruption of their homeostatic functions, increased expression of interferon and inflammatory pathway-associated genes ^27^. *Btg2* (B-cell translocation gene 2) has been reported to negatively regulate the proliferation of glial cells in response to cerebral hypoperfusion in animal cerebral ischemia models ^28^. *Txnip* (thioredoxin interacting protein), as an endogenous thioredoxin inhibitor, inhibits the antioxidant effects of thioredoxin, which have been found to mediate the inflammatory signal of NLRP3 activated by glucocorticoid in mouse microglia ^29^. *Hspa1a (hsp70)* has been identified as the ligand of toll-like receptors TLR2 and TLR4 ^30^, whose activation has been investigated for therapeutic potential in neurological diseases ^31, 32^. A clinical study reported that the upregulation of *Malat1* in cerebrospinal fluid (CSF) could accurately predict cerebral vasospasm in SAH patients ^33^. Collectively, with function pathway analysis, the SMG-C4 highly expressed genes were found related to leukocyte activation, positive regulation of cellular metabolic process and positive regulation of nitrogen compound metabolic process (**Figure S7-B and Figure S10-B**).

#### 3.2.2. Activated microglia: SMG-C2

SMG-C2 was characterized by high expression levels of *Ctsb*, *Ctsd*, *Ccl3*, *Ccl4, Lgals3bp*, *Cst7, Grn* and *Nfkbia etc.*. High *Ctsb* and *Ctsd* expression implicates elevated lysosomal activity in this MG cluster, which might be linked to enhanced phagocytic capacity ^34^. *Lgals3bp* encodes the galectin 3–binding protein, which is a secretory glycoprotein. Prior studies showed that the *Lgals3* (Galectin 3] gene is usually expressed in early postnatal brain regions and otherwise almost absent in adulthood ^34, 35^. SMG-C2 also showed elevated expression of *Serpine1*, encoding PAI-1– a regulator of tissue plasminogen activator that has been found connected with the prognosis of SAH patients ^36^. Notably, *Ctsb*, *Ctsd* and *Cst7* were also upregulated in the disease-associated microglial (DAM) identified in the 5xFAD mouse model of Alzheimer’s disease ^37^. Together with the elevated expression of cytokine-encoding genes *Ccl3* and *Ccl4*, these features suggested that SMG-C2 represents activated microglia. Pathway enrichment analysis showed that SMG-C2 transcriptomes were enriched in genes involved in regulating metal ion transport, monocyte chemotaxis, regulation of inflammatory response and regulation of hydrolase activity (**Figure S6-C and Figure S9-C**).

#### 3.2.3. SAH-associated microglia (SAM): SMG-C5

Among the MG subclusters, we identified a subcluster (SMG-C5) that partially resembles DAM. SMG-C5 was characterized by the expression of marker genes, including *Spp1, Lpl, Apoe, Ctsb, Lgals1, Lgals3, Fabp5, Mif, Lilrb4a, Lyz2, CD63, Cst7* and *Vim* **Figure 2**, **Figure 4 and Excel S2**). Other genes with high expression in (SMG-C5 included *Anxa2, Apoc2, CD63, CD72, Ctsc, Ctsz* and *Ctsl etc.* – which have also been found upregulated in developing mouse brain microglia, AD and lysolecithin (LPC) injury mouse models ^37–41^. *Spp1* (encoding secreted phosphoprotein 1, also known as osteopontin) has the potential to enhance myelination. In the developing mouse brain, the axon tract-associated microglia (ATM) are amoeboid-shaped *Spp1+* microglia with upregulated expression of *Lgals1* (encoding galectin-1) and *Lgals3* (encoding galectin-3) ^38^. *Apoe* (encoding apolipoprotein E), an AD-risk gene, is also known to regulate the reactivity of microglia ^42^. It has been found that the ApoE4 genotype negatively affects cognitive morbidity and delayed ischemic neurologic deficit recovery in SAH patients ^43^. *Mif* (macrophage migration inhibitory factor) and *Fabp5* are related to cell growth, motility, inflammation and immunomodulation in macrophages and other cells ^44–46^. *Lilrb4a* and *Lyz2* were shown as phagocytic biomarkers in adult cerebellar microglia ^38^ (although we excluded the cerebellum from the brain tissue harvesting step). *Lpl* (encoding lipoprotein lipase) is connected with the metabolism and phagocytic capacity of microglia. Evidence also suggested that *Lpl* is a feature of reparative microglia that could regulate remyelination and repair through the clearance of lipid debris ^47, 48^.

**Figure legend 2:**
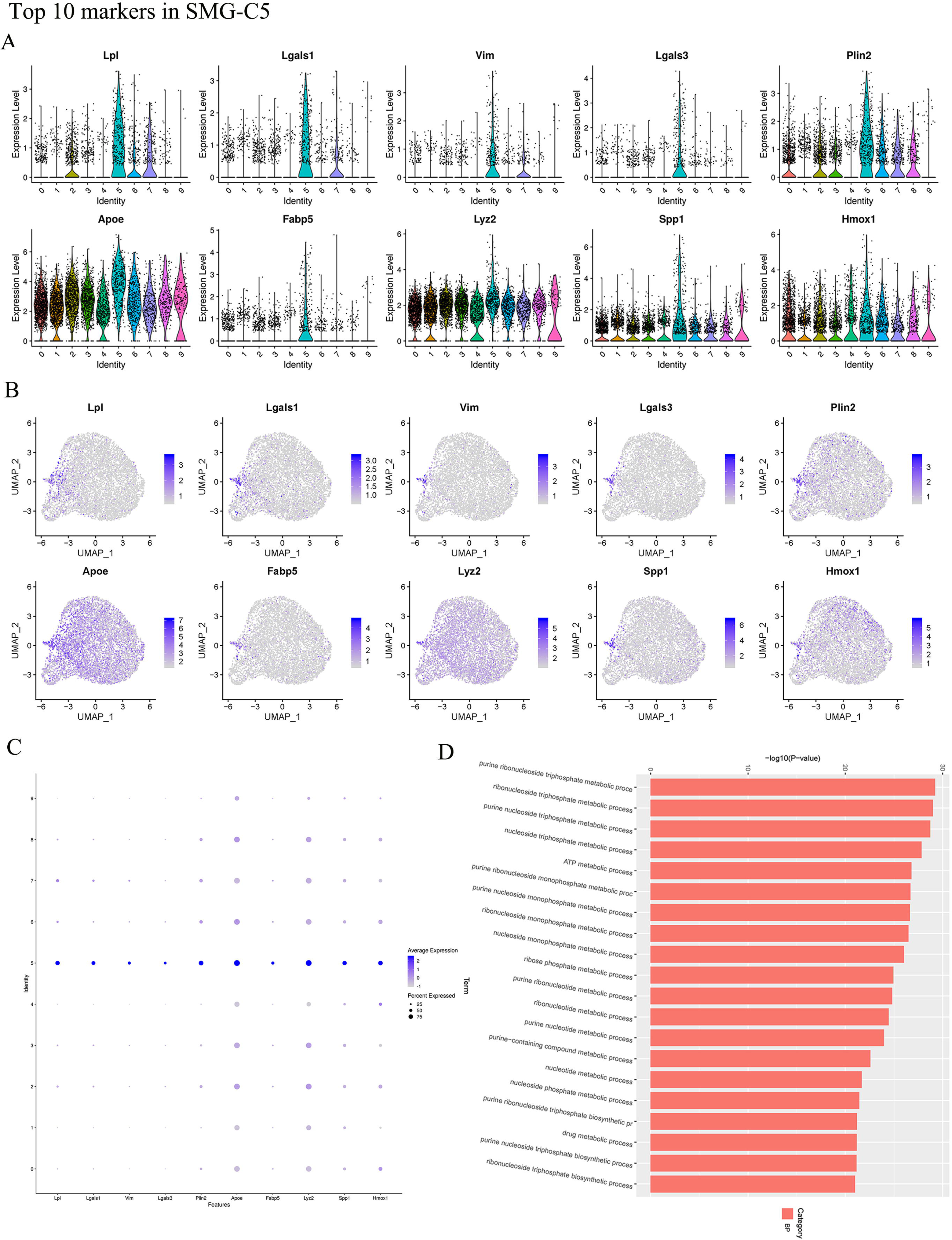
A: Violin atlas of top 10 genes in SAH microglia cluster 5 (SMG-C5); B: Top 10 genes distribution in SMG-C5; C: Top 10 genes expression dot plot in SMG-C5; D: Enrichment of SMG-C5.

**Figure legend 3:**
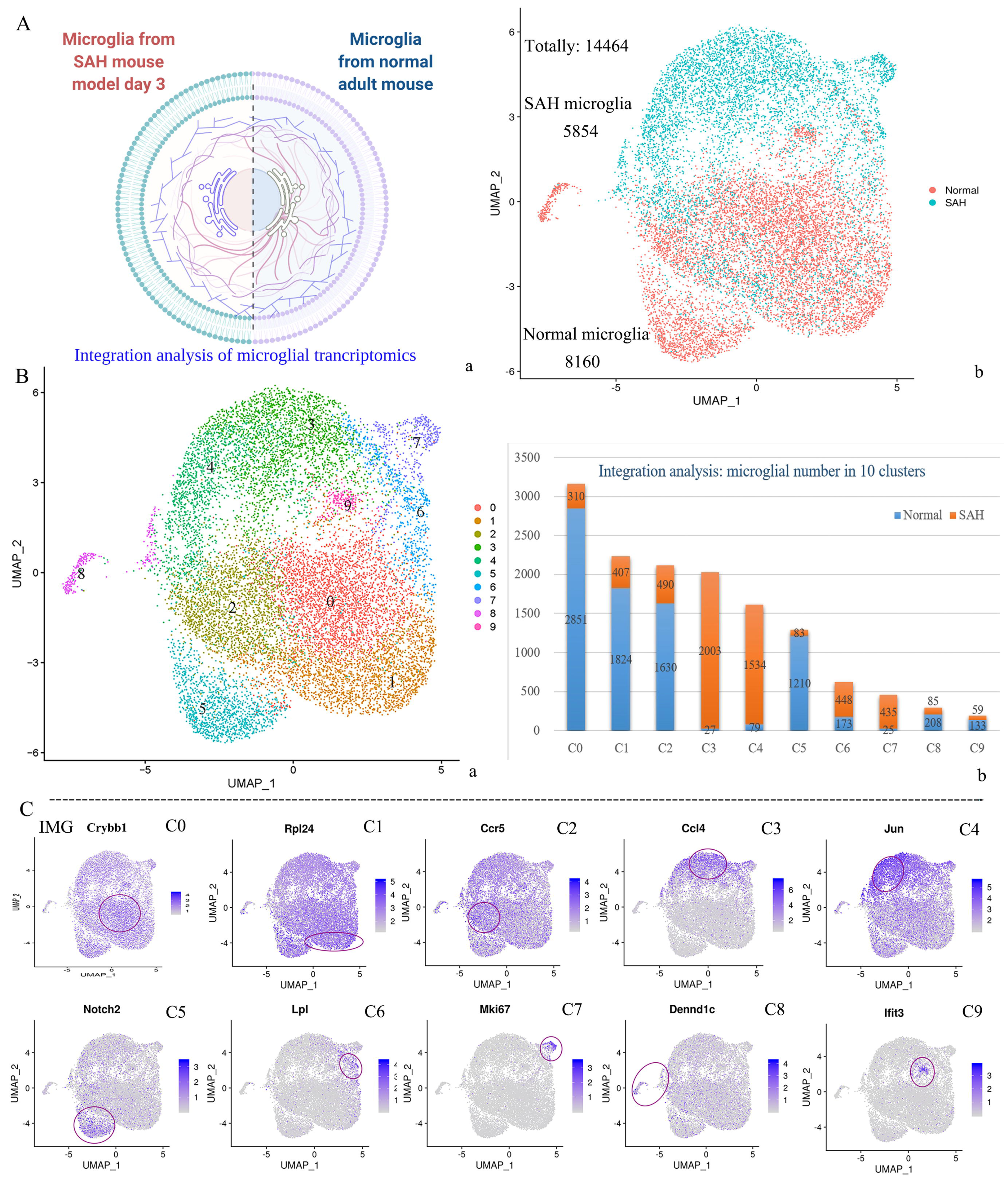
A: a-Mapping of SAH and normal microglia samples; B: A-UMAP of Integration analysis of SAH and normal microglia (IMG), b-SAH and normal microglia numbers in each IMG cluster; C: Biomarkers of each clusters in IMG.

Based on prior reports, DAM found in AD, injury responsive microglia (IRM) found in LPC injury and ATM found in the developing mouse brain all share the expression of a core set of genes encompassing *Spp1*, *Apoe* and *Lpl* ^38^. In SAM, these genes are all upregulated. On the other hand, important differences in the transcriptomes of SMG-C5 vs. DAM indicates that microglia have a distinct way of activation in SAH. For example, unlike DAM, SAM does not have elevated expression of *Trem2, Clec7a, Ccl6, or Axl*, but SAM has its unique panel of highly expressed genes, such as *Spp1 Lgals1, Lgals3, Fabp5, Mif, Lilrb4a* and *Vim etc.*^37^

Collectively, based on these transcriptomic features we termed microglia of SMG-C5 the SAH-associated microglia (SAM), to emphasize the cluster’s special activated state post-SAH. From gene ontology (GO) and Kyoto Encyclopedia of Genes and Genomes (KEGG) enrichment analysis, we found that the upregulated genes in SAM are related to oxidative phosphorylation (gene number: 31, p = 5.42 × 10^-27^), lysosome (gene number: 10, p < 0.01), apoptosis (gene number: 10, p < 0.01), glycolysis/gluconeogenesis (gene number: 11, p = 3.68 × 10^-07^) (**Figure 5-A**).

**Figure legend 4:**
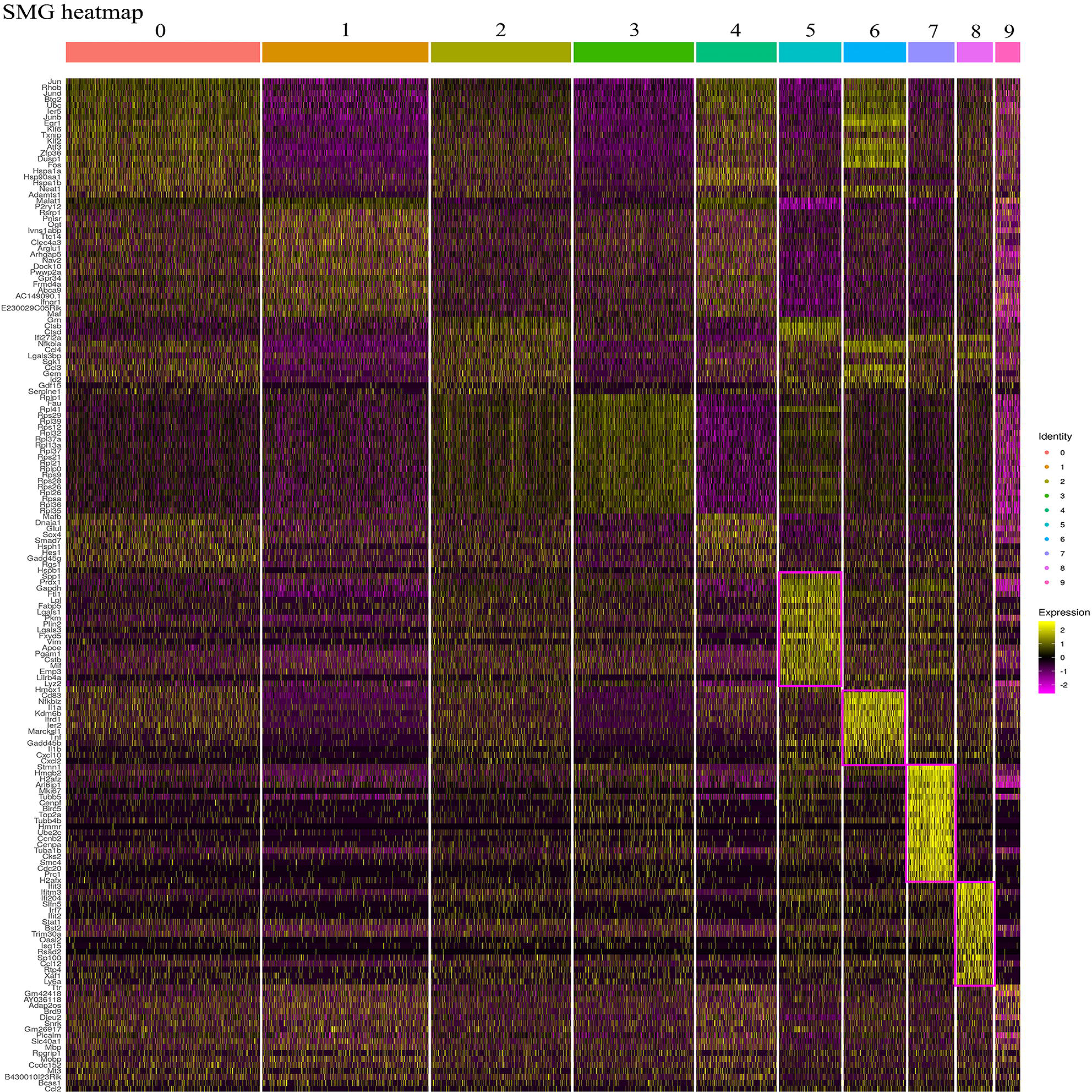
Gene expression of SAH microglia (SMG-heatmap).

**Figure legend 5:**
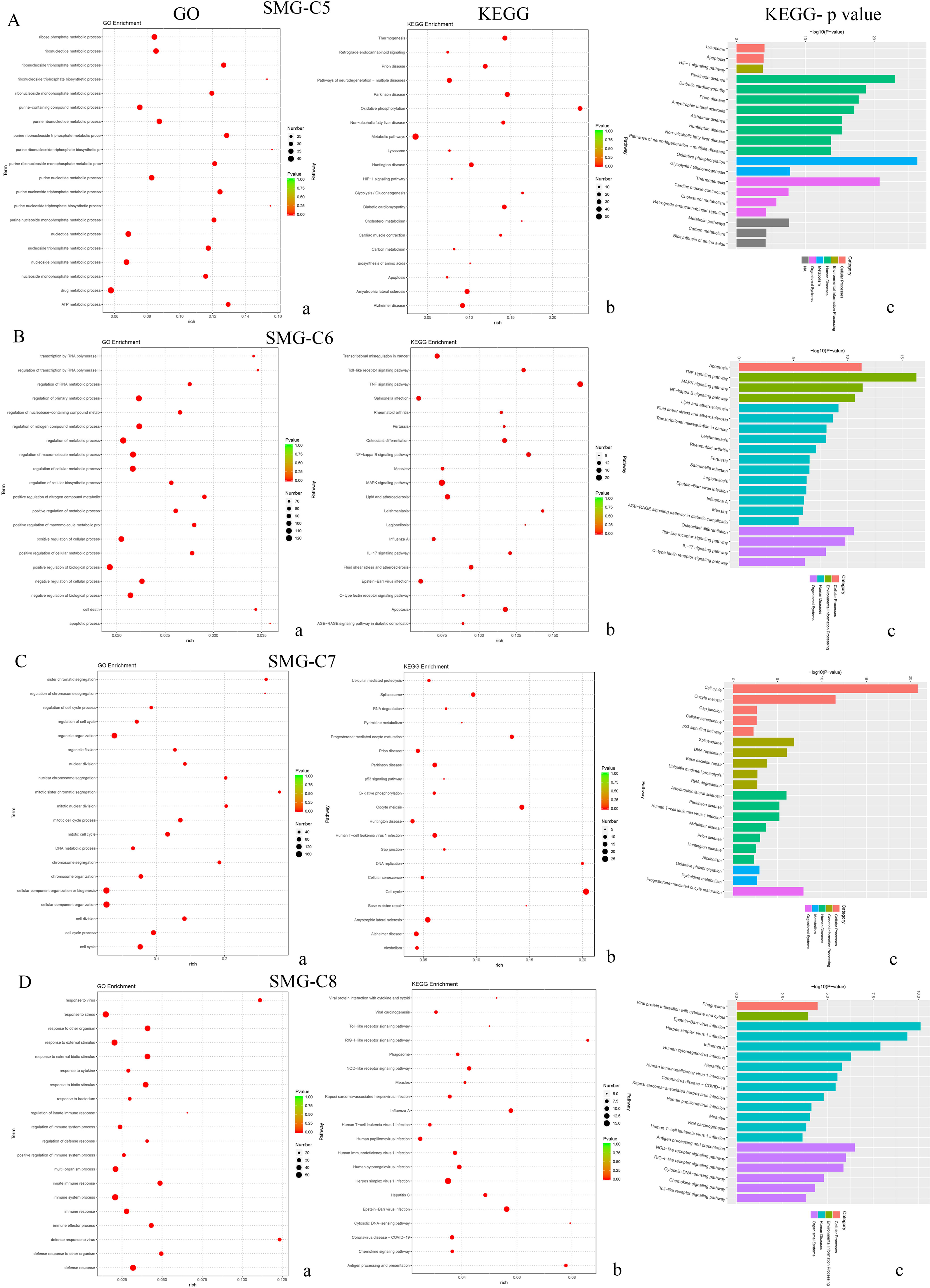
Enrichment analysis of SAH microglia (SMG). A: SMG-C5; B: SMG-C6; C: SMG-C7 and D: SMG-C8. a: GO, b-c: KEGG.

#### 3.2.4. Inflammation-associated microglia (IAM): SMG-C6

The gene expression pattern in SMG-C6 suggests that it represents a class of inflammation-associated microglia (IAM) in SAH (**Excel S2, Figure 4 and Figure S8-A**). SMG-C6 microglia expressed a variety of cytokines (included *Il1a* (encoding interleukin 1 alpha), *Il1b* (encoding interleukin 1 beta)*, Tnf* (encoding tumor necrosis factor α), *Ccl4* and *Ccl3*), chemokines (including *Cxcl10* (C-X-C Motif Chemokine Ligand 10), *Cxcl2* (C-X-C Motif Chemokine Ligand 2)), and other immune-signalling regulatory genes (including *CD83, CD74* (MHC class II), *CD14*, *Nfkbia* and *Nfkbiz* (NF-κB inhibitor Zeta)). Overexpression of *Il1a, Ccl4* and *Ccl3* has been observed in the IRM subpopulation in multiple sclerosis (MS) mouse model, where *Ccl4* was also a marker in human MS ^38, 39^. Over-activation of IL-1β-mediated signalling has been reported to interrupt the dendritic spine formation and interfere with memory consolidation ^26^. *Cxcl10* and *Cxcl2* are interferon response genes, which are also highly expressed in the experimental autoimmune encephalomyelitis (EAE) mouse model of MS ^49^. Results of pathway enrichment analysis suggested that SMG-C6-enriched genes are related to positive regulation of macromolecule metabolic process, regulation of the primary metabolic process, and positive regulation of the cellular metabolic process. Consistent with the putative IAM state inferred from marker gene expression patterns, KEGG results also suggested that SMG-C6 were enriched in genes related to interleukin, tumor necrosis factor (TNF), toll-like receptor 4 (TLR4) and nuclear factor-kappa B (NF-κB) signaling pathways (**Figure 5-B**). Indeed, these signaling pathways have been found to participate in SAH-induced neuroinflammation and affect SAH outcomes ^50–52^.

#### 3.2.5. Proliferation-associated microglia (PAM): SMG-C7, and interferon responsive microglia: SMG-C8

SMG-C7 have high expression of proliferative and cell cycle genes (e.g., *Birc5* (baculoviral IAP repeat-containing 5), *Ccnb2*, *Cenpa, Cenpf, Mcm5, Ube2c, H2afz*, *H2afx, Cdks and Mki67*) (**Figure 4**, **Figure S8-B and Excel S2**), which are usually expressed in developing but not adult mouse microglia ^53^. These suggested possible reactivation of developmental pathways in SAH ^38^, and we thus termed SMG-C7 microglia proliferation-associate microglia (PAM). In line with this, pathway enrichment results indicated that SMG-C7 was enriched with genes associated with mitotic nuclear division, cell division and cell cycle (**Figure 5-C**). SMG-C8 marker genes were characterized by several interferon responsive genes (including *Ifit3, Ifitm3, Ifi204, Slfn5, Irf7, Ifit2, Ifi27l2a* and *Cxcl10*). SMG-C8 also has a high expression of MHC class I related genes (including *H2-D1, H2-K1, H2-T23* and *H2-Q7*) ^54^. In GO analysis, SMG-C8 was suggested to be highly responsive to cytokines and other biological stimuli, potentially mediated by the MHC class I protein complex, chemokine activity regulation and CCR chemokine receptor binding (**Figure 5-D**). Finally, SMG-C9 abundantly expressed *Ttr, Mbp, Mobp, Myh9, Slc40a1* and *Dennd1c*. These genes may be connected with myelin and oligodendrocyte function ^55^. Furthermore, SMG-C9 was enriched with genes related to the regulation of the metabolism, RNA metabolic processes, lysosomes and ABC transporters (**Figure S7-C**).

### 3.3. Significant upregulated signaling pathways in SMG sub-populations’ interaction (Microglia subsets interaction)

In all, the above results revealed the wide range of microglia states in SAH. However, microglia subtypes do not act in isolation, and in general all brain cell types closely interact to mediate diverse physiological and pathological processes. We thus then carried out cell-cell communication analysis.

In total, 15 important intracellular signaling pathways related to neuroinflammation were detected (**Figure 6**). Significantly up-regulated signal pathways found, according to the order of expression, are as follow: chemokine ligands (CCL), GALECTIN, transforming growth factor beta (TGF-β), junction adhesion molecule (JAM), intercellular adhesion molecule (ICAM), growth arrest-specific (GAS), secreted phosphoprotein 1 (SPP1), progranulin (GRN), semaphorin 4 (SEMA4), cell adhesion molecule (CADM), colony-stimulating factor (CSF), TNF, amyloid-β precursor protein (APP), growth differentiation factor (GDF) and oncostatin M (OSM). Specific microglia clusters participated in each signal pathway (see the detailed relationships between each signal pathway and SMG clusters summarized in **Figure 7**, **Figure S11**, **Figure S12 and Figure S13**). TGF-β could promote human leptomeningeal cell proliferation within the first 48 to 72 hours after SAH, and the intracerebral injection of TGF-β caused hydrocephalus to develop in mice models ^56, 57^.GALECTIN was associated with injury and neuroinflammation after SAH, animal studies found Inhibiting Galectin-3 could prevent blood-brain barrier disruption in SAH ^58^, clinical studies found the increased level of galectin-3 in SAH patients is related to poor outcome ^52^.

**Figure legend 6:**
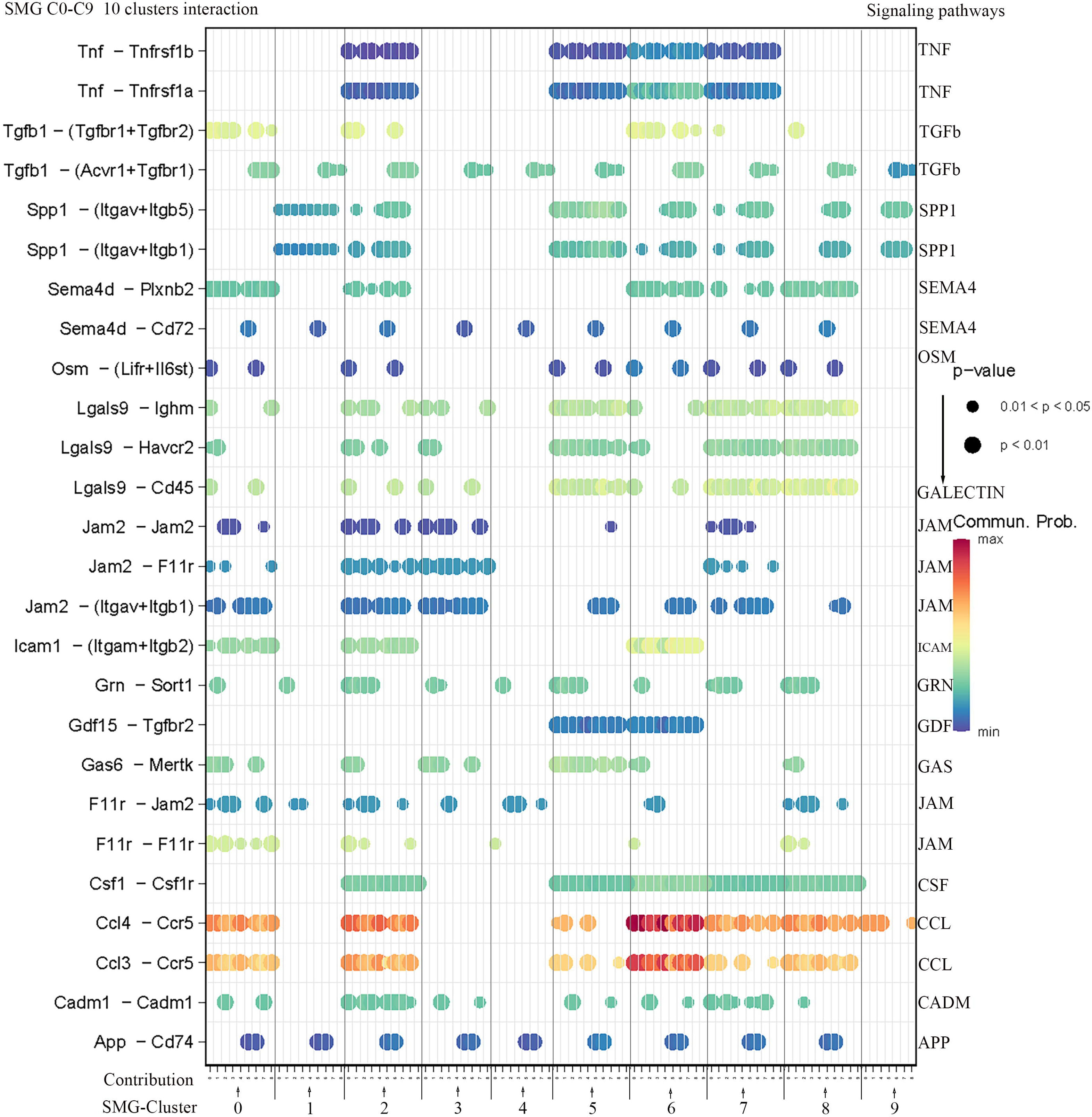
SMG microglia subsets interaction. (Totally 15 signal pathways.)

**Figure legend 7:**
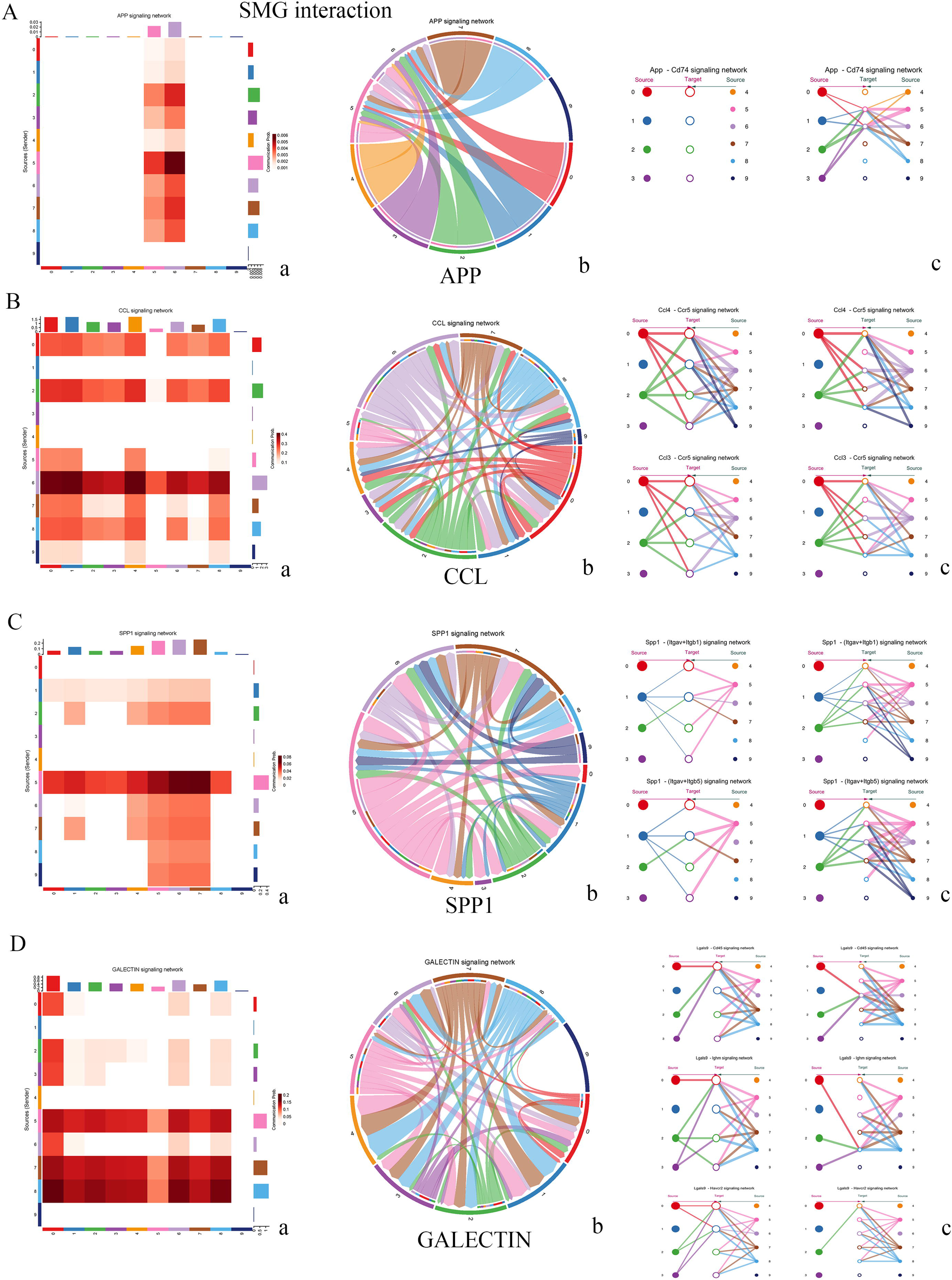
SMG microglia subsets interaction of the specific signaling pathways. A: APP; B: CCL; C: SPP1 and D: GALECTIN signaling pathway. a: heatmap, b: chord map and c: hierarchy connection.

SMG-C5 and SMG-C6 appeared to be the main targets of the APP signal pathway (**Figure 7-A**), suggesting a potential connection between SAH prognosis and APP activation ^42^. SMG-C5 (SAM) and SMG-C6 (IAM) are also important putative participants in CCL, GALECTIN, TGFb, SPP1, and TNF-mediated signaling pathways. Collectively, these results indicated that microglia as a whole participate in the inflammatory reaction of the brain after SAH, but specific subpopulation participates differentially in mediating neuroinflammatory responses and pointed out the necessity of a microglial subgroup-based analysis strategy. Overall, SMG-C5, SMG-C6 and SMG-C7 have the most interaction with other clusters, and warrant further in vivo studies to elucidate their functional roles in SAH (**Figure 7**, **Figure S11**, **Figure S12 and Figure S13**).

### 3.4. Intercellular communication analysis revealed the important roles of microglia after SAH and the complexity of neuroinflammation

Early brain injury (EBI) is one of the major causes of mortality and morbidity in SAH patients. The pathophysiological mechanisms of EBI are complex and not fully understood. We compared the gene expression of the extracted microglia with that of other kinds of cell (**Excel S4**), and found multiple pathways enriched specifically in microglia, including leukocyte activation, migration and adhesion, glia cell migration and activation, gliogenesis, regulation of inflammatory response, lysosomal pathways (**Figure S18**). In addition, apoptosis, p53-related, mitochondria and caspase signaling pathways, which were found upregulated in microglia, has also been found connected with EBI after SAH in previous evidence ^9^ (**Figure S18**).

As microglia also closely interact with other brain cell types, we also performed inter-cell type communications between microglia and other cells (**Figure 8**, **Figure 9**, **Figure S14, S15, S16, S17 and S18**). Microglia have been found to participate in most post-SAH inflammatory pathways (25 signaling pathways in total), among which 13 signaling pathways involve interactions with neurons (including GAS, JAM, CHEMERIN, CSF, PSAP, PTN and SPP1-mediated pathways) (**Figure S14 and Figure S16**). Microglia, CNS-associated macrophage, macrophage and astrocyte also closely interact in the post-SAH inflammation (**Figure 8**, pattern 2), via CCL, galectin, PSAP, PROS, GRN, CD45, CD39 and TGF-β (outgoing), and CCL, TGF-β, galectin, GAS, ICAM and JAM (incoming) signaling (**Figure 8**). CCL, galectin and TGF-β are contributed by several genes and cell types (**Figure 9**), while they are also highly implicated in microglial subsets interaction (**Figure 6**).

**Figure legend 8:**
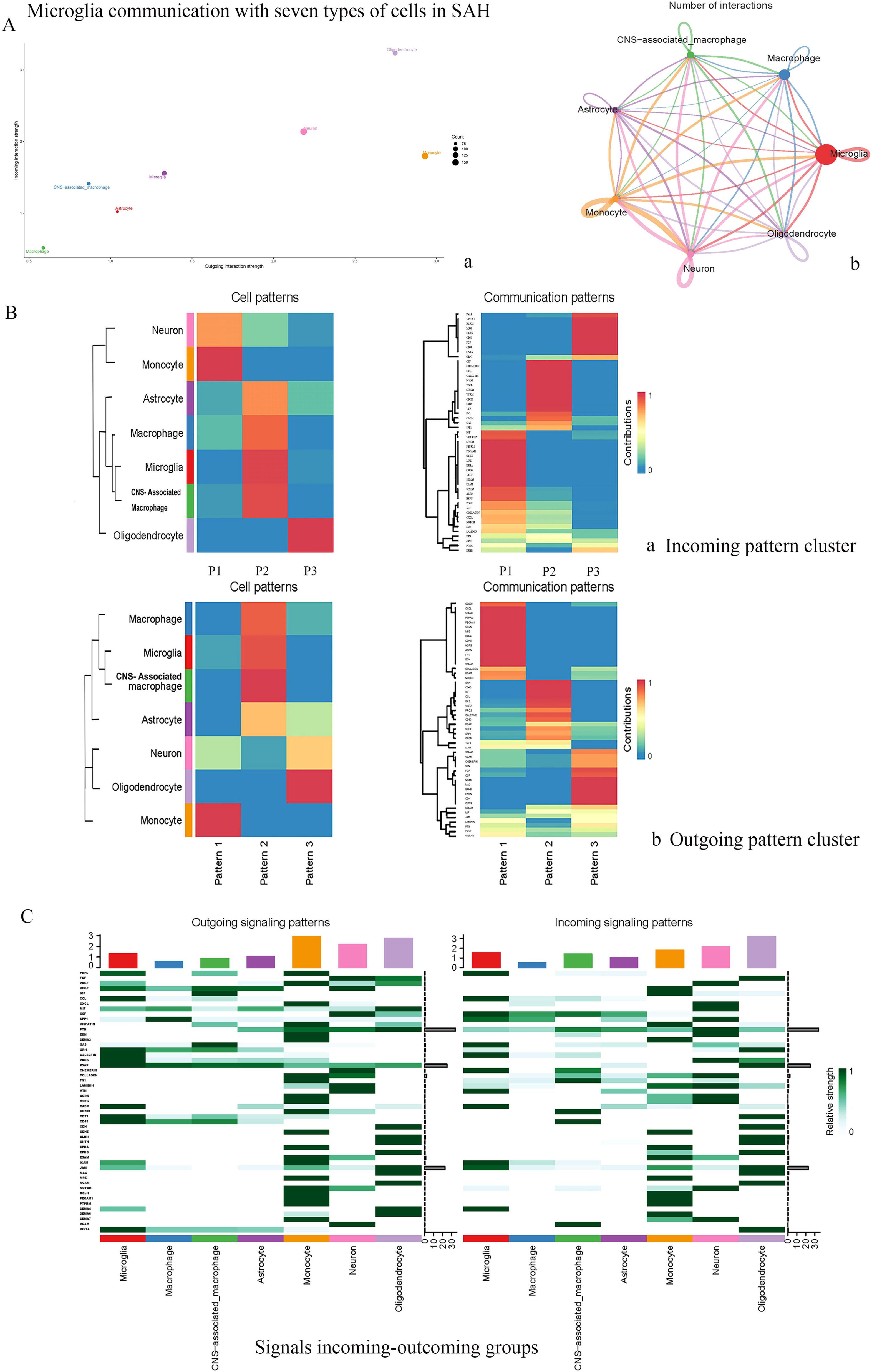
Microglia interactions with seven types of cells. A: a: Cell communicating strength, b: cell interactions; B: a: Cell incoming signal patterns and b: cell outgoing signal patterns; C: Signals incoming and outgoing groups.

**Figure legend 9:**
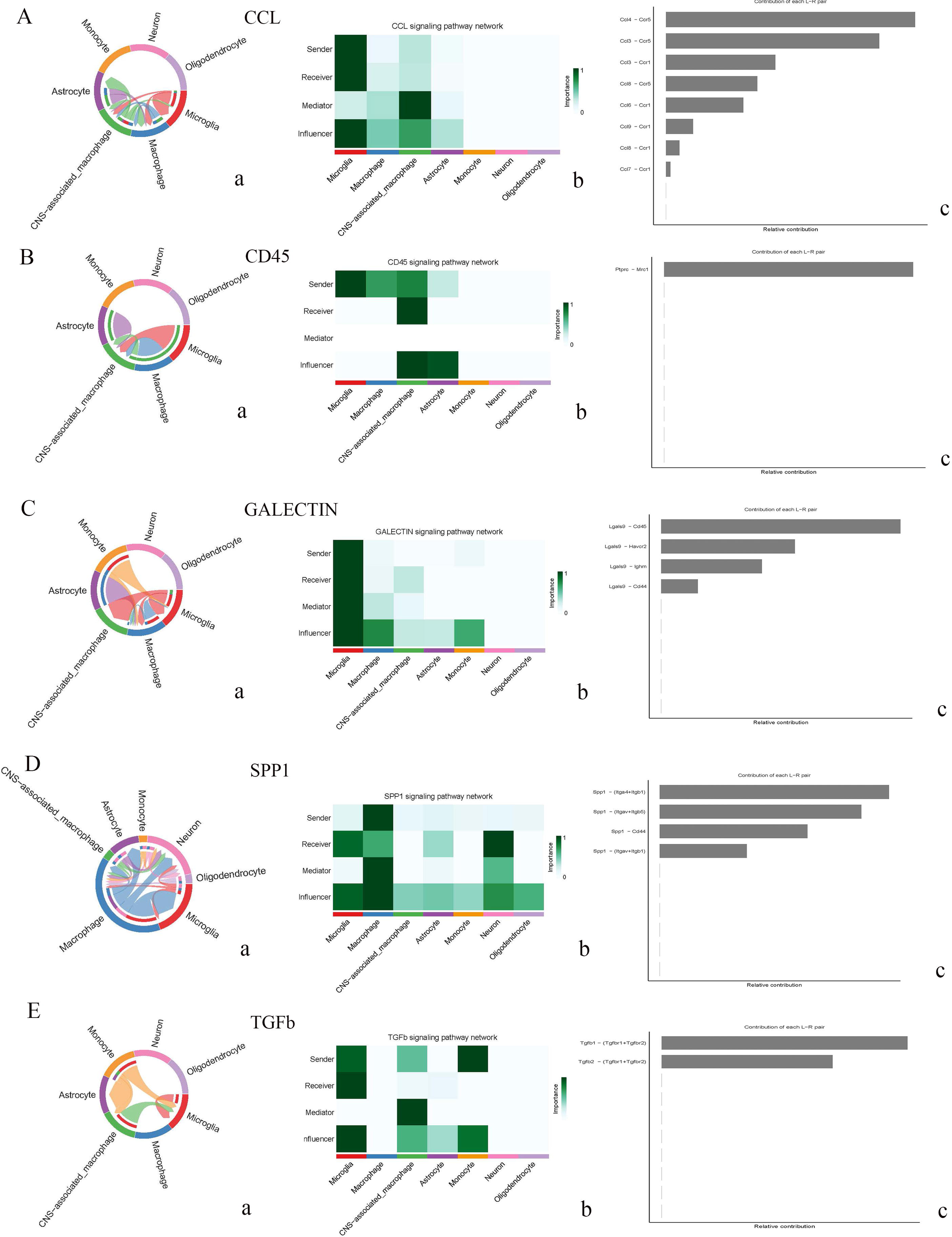
Signaling pathways expressions-microglia interacts with seven types of cells. A: CCL; B: CD45; C: GALECTIN; D: SPP1 and E: TGFb. a: chord map, b: contributing role and c: signaling role hierarchy.

Together, these results suggested that microglia likely play an important role in EBI, while other types of brain cells may also play significant complementary roles in a coordinated manner with microglia ^59^.

### 3.5. Trajectory state analysis indicating the transcriptional state complexity of SMG subpopulation

We then performed branch expression analysis modeling (BEAM) on the SMG transcriptomes. SAH microglia were divided into 5 states (state 1 to state 5) in pseudotime analysis (**Figure 10-A**). With monocle analysis, we noted that all subpopulations contain a variety of cell states. SMG-C4 and SMG-C9 were probably in an intermediate stage because they mainly contained states 1 and 4, but not state 5.

**Figure legend 10:**
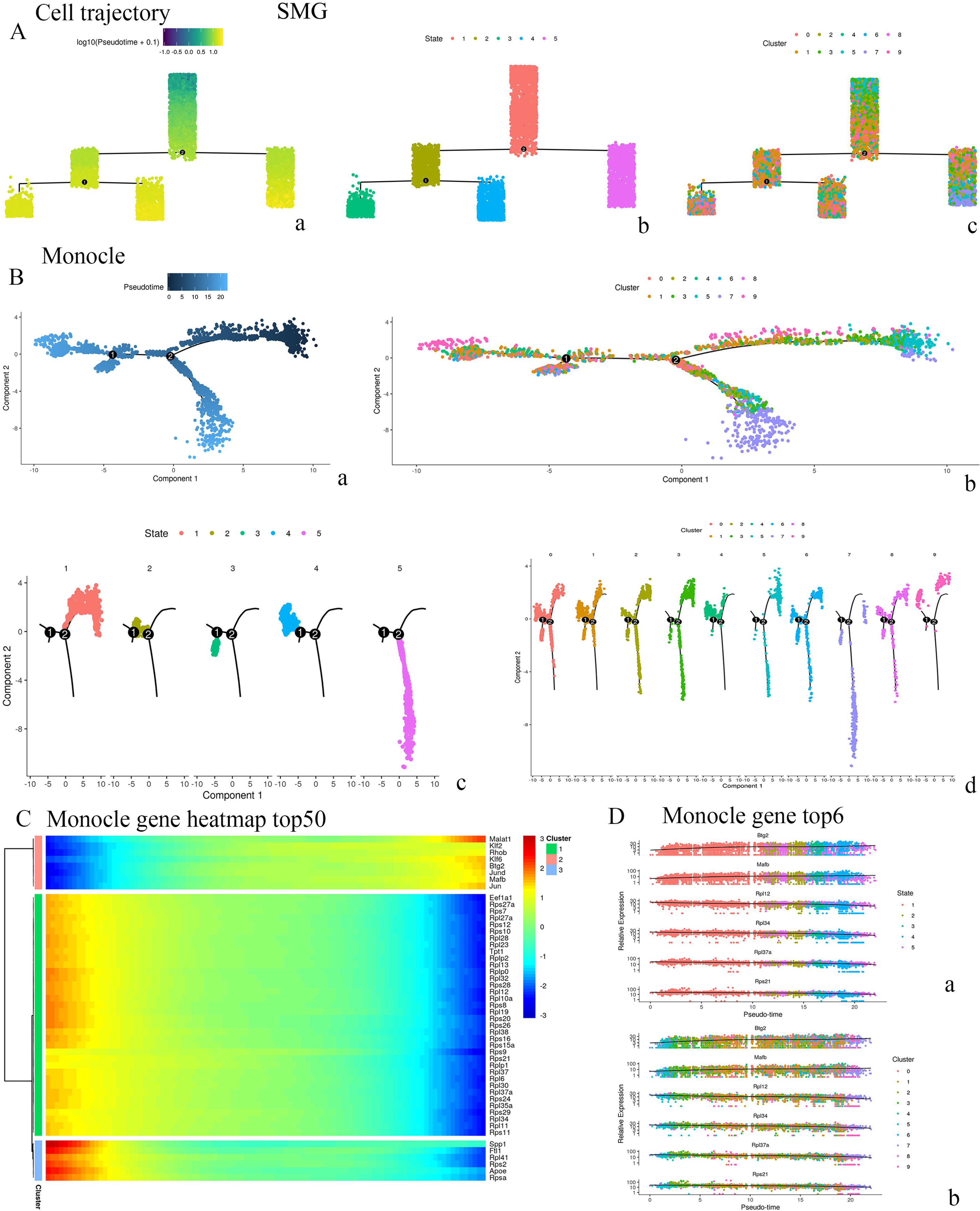
Trajectory analysis of SMG (10 clusters). A: Microglia trajectory; B: Monocle branch (a: Pseudotime, b: monocle branch (cluster), c: monocle branch (state), d: monocle branch (cluster split)); C: monocle genes heatmap; D: monocle genes state.

The entire process of 5 states can be found in SMG-C2, SMG-C6 and SMG-C8. SMG-C5, SMG-C6, SMG-C7 and SMG-C8 included the late states of microglia transformation (**Figure 10-B**). (**Figure 10-B**).

Among the top 6 genes that varied with pseudotime, *Btg2* and *Mafb* gradually become unregulated and *Rpl12, Rpl34, Rpl37a* and *Rps21* were progressively downregulated from states 1 to 5 (**Figure 10-C**). From the heatmap of top 50 genes with pseudotime, the immediate early genes (*Jun, Jund, Klf6, Klf2 -* enriched in SMG-C0), transcriptional regulatory genes (*Btg2, Rhob* and *Mafb -* enriched in SMG-C4) and *Malat1* were all upregulated with pseudotime (**Figure 10-C**), which indicates that SMG-C0 and SMG-C4 represented important states in microglial activation and transition. At the same time, *Spp1, Apoe* and *Ftl1* (enriched in SAM), ribosome genes (*Rps41, Rps2* and *Rpsa*), are downregulated from state 1 to 5. These suggested that SAM and ribosomal genes became upregulated early after SAH, and that SAM may appear earlier than IAM and PAM based on the trajectory analysis. At the cellular level, the overexpression of *Rps4l* could weaken cell viability and proliferation, and suppress cell cycle progression ^60^. The expression of *Rps2* has been reported to be elevated in SAH patients’ blood ^61^, and the up-regulated expression of *Malat1* in CSF of SAH patients was also related to cerebral vasospasm and poor outcome ^33^. These results showed that SAM appeared earlier than IAM and PAM in the trajectory analysis. The transformation of microglia may be closely related to these genes.

### 3.6. Integration analysis of the single-cell transcriptome of SAH and normal microglia

We then performed comparative analysis of SAH (*n* = 5,854) and normal (*n* = 8,160) mouse brain microglia, to further reveal their transcriptomic differences (**Figure 3-A**). All the microglia transcriptomes were similarly clustered based on their gene expression characteristics (**Excel S3, Figure S19**), and termed integrated microglia (IMG) clusters 01-09 (IMG-C0 to 09). SAH and normal mouse brain microglia had different proportions and markers among these ten clusters (**Figure 3-B, C**). We compared the gene expression characteristics between IMG and SMG clusters and found some similar clusters corresponding to SAM, IAM and PAM (**Table 1).** The upregulated and downregulated genes in SAH microglia were also calculated and compared in each IMG group (**Table 2**).

**Table 1:**
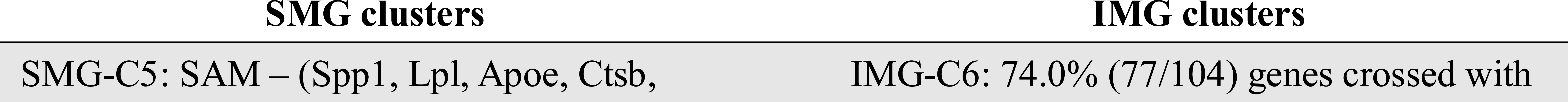
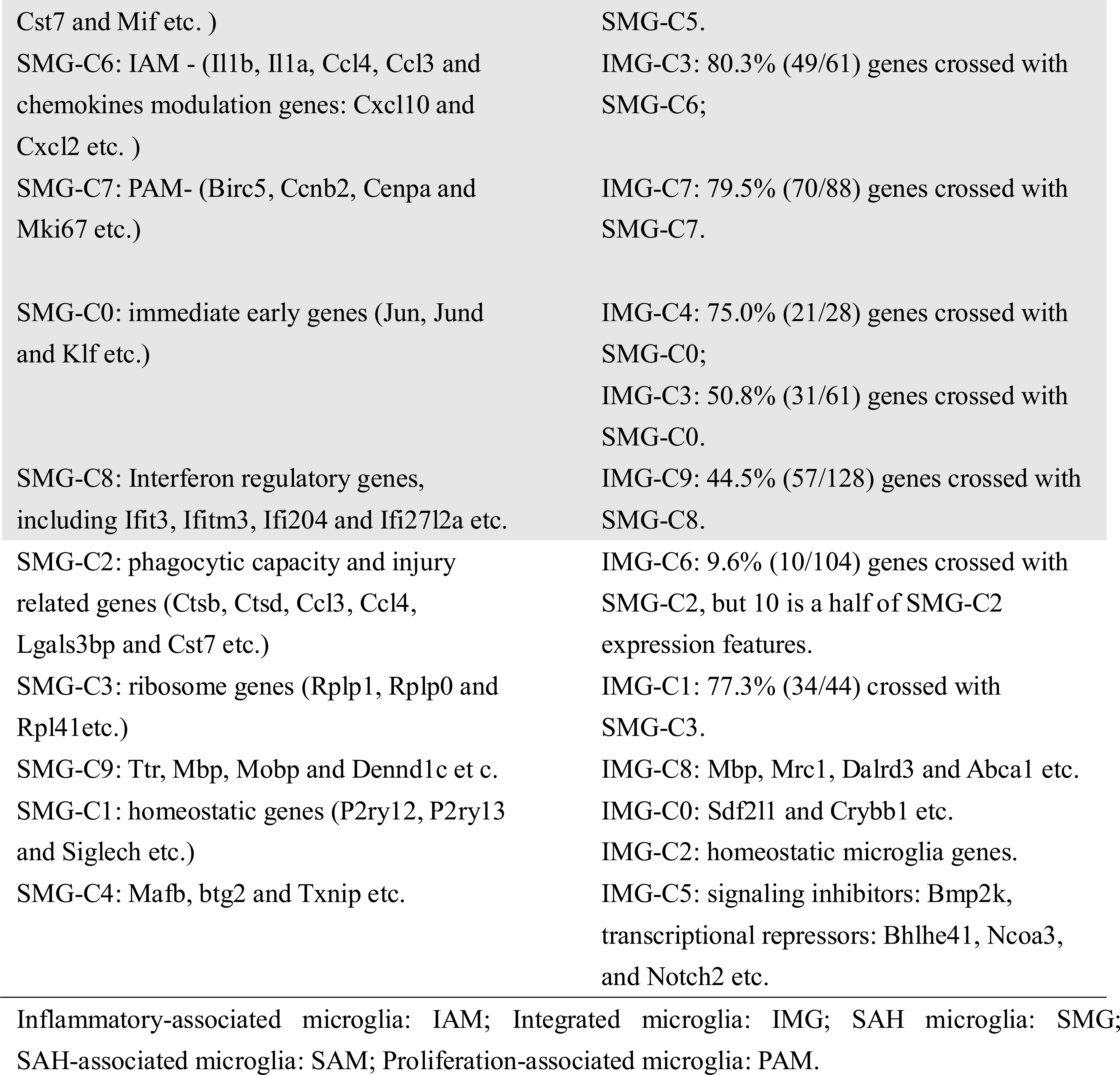
Gene comparison between IMG and SMG clusters.

**Table 2:**
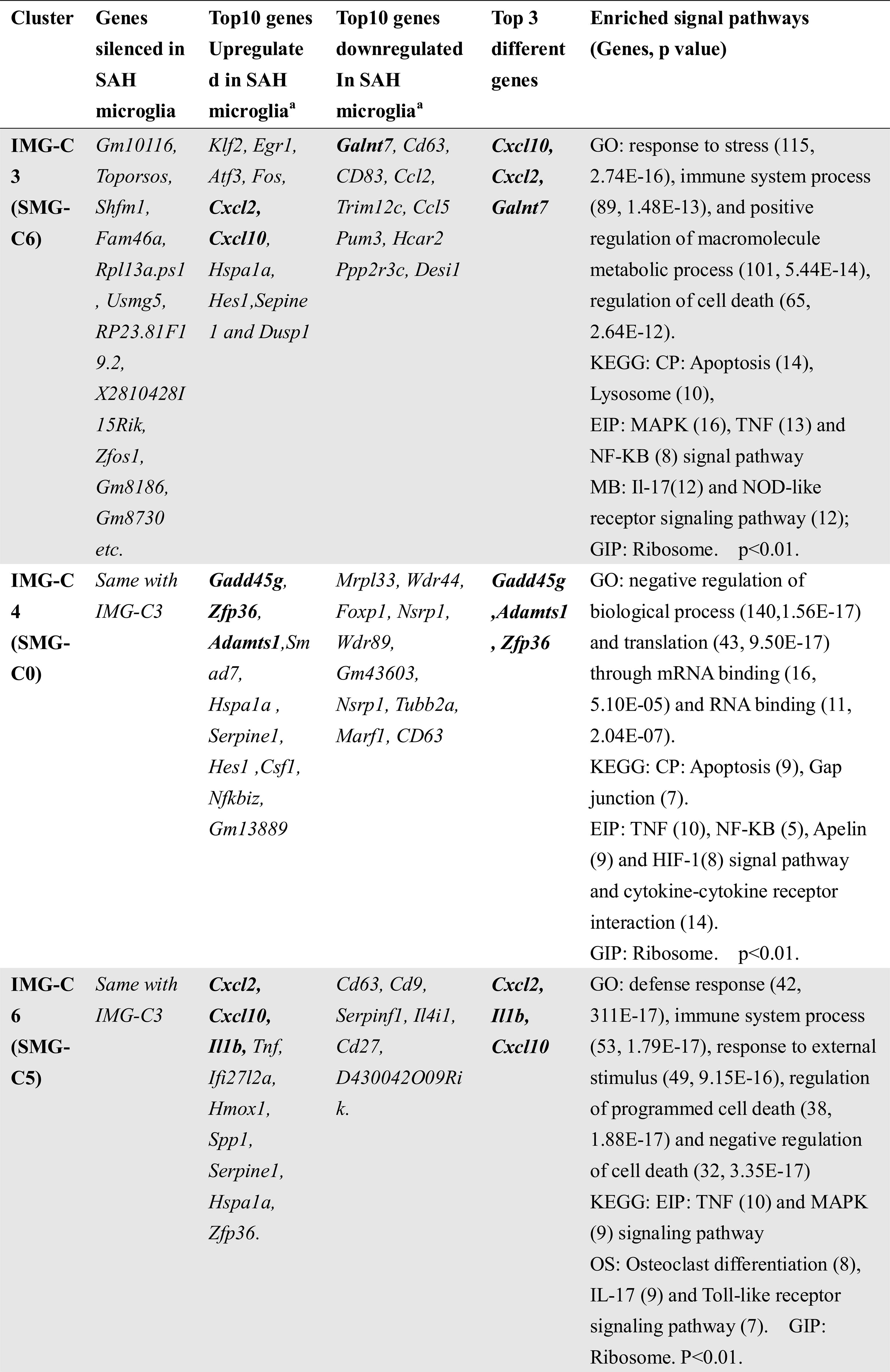
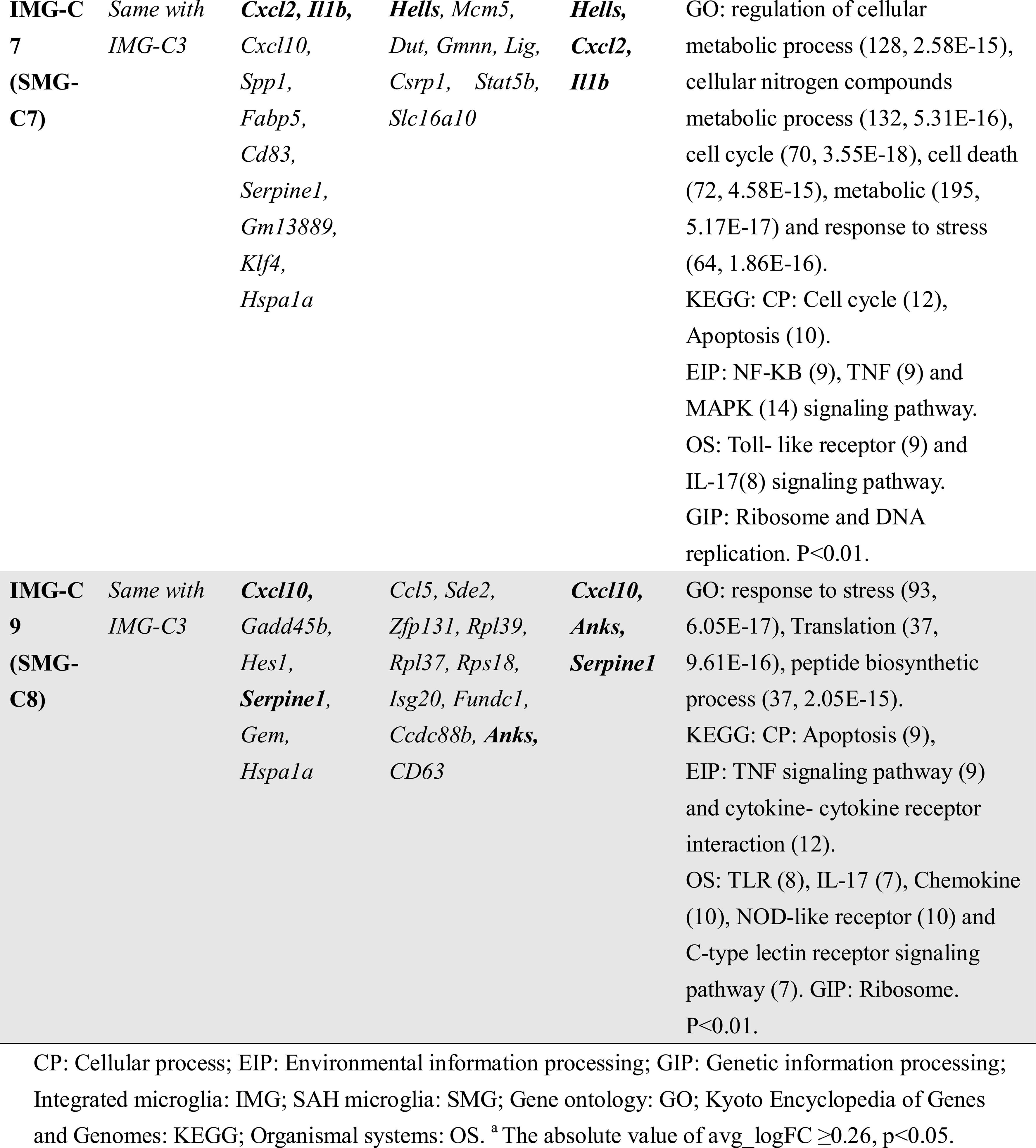
Different gene expression of SMG and normal microglia in IMG clusters.

### 3.7. Transcriptome difference of SAM, IAM and PAM with normal microglia

From the IMG clustering results, most of the normal mice microglia were classified as IMG-C0 (*Sdf2l1* and *Crybb1)* ^62^, C1 (ribosome genes), C2 (*Ccr5 and Zbtb20 etc.*) and C5 (signaling inhibitors: *Bmp2k,* transcriptional repressor: *Bhlhe41, Ncoa3*, and *Notch2 etc.*), while the other IMG clusters were predominantly from SAH microglia **(Excel S3, Figure S19**). IMG-C6 (74.0% genes), IMG-C3 (80.3% genes), and IMG-C7 (79.5% genes) were clusters with gene markers resembling SMG-C5 (DAM cluster), SMG-C6 (IAM cluster), and SMG-C7 (PAM cluster), respectively (Marker gene were compared) (**Table 1**).

Compared with normal microglia in each IMG clusters, SAH microglia in IMG-C6, IMG-C7 and IMG-C3 exhibited upregulated expression of *Cxcl2, Cxcl10, Il1b, Hmox1, Spp1, Apoe, Lpl, Cd83*, *Serpine1, Dusp1* and *Hes1 etc.* which are the most distinct genes for DAM, IAM and PAM, may connected with the activation of these clusters. (**Table 2**) These differentially expressed genes provided hints on the possible mechanism of SAM, IAM and PAM generation. *Cxcl2* has been observed upregulated in the cerebral arteries of SAH animal models ^63^. Inhabiting microglia specific *Cxcl2* expression could alleviate ischemic brain injury by reducing neutrophil infiltration in ischemic stroke ^64^. *Hmox1*-encoded heme oxygenase-1 has been found to have an anti-inflammatory effect when activated in lipopolysaccharide-activated microglia, and confers neuroprotection in ischemia stroke ^65, 66^. Most importantly, a study found that the elimination of *Hmox1* from microglia would damage the uptake of extravasated erythrocytes, aggravate the neuronal cell death in experimental SAH, which is very important for the recovery of aneurysmal SAH patients ^67^. Evidence showed that *IL-1b* could be an important modulator of microglial activation and proliferation, studies have found that the production of Il-1b in microglia is P2X7R pore-dependent ^68^. In previous study, *Cst7, Cd9* and *Cd63* were over expressed in DAM microglia of neurodegenerative disease ^37^, but here *CD9* and *CD63* were low expressed in IMG-C6 (SAM). Microglia with different expressions of *Cd83* and *Cd63* were related to specific functional changes of microglia sub-states ^69, 70^, and here both downregulated in SAM and IAM.

*Serpine1* and *Hspa1a* were upregulated in almost all SAH microglia of IMG subsets. Clinical evidence suggested an important regulatory role of the *Serpine1* gene polymorphism in poor clinical outcomes of SAH ^36^. *Serpine1* could also regulate the inflammatory response of microglia during brain inflammation ^71^. Heat shock proteins and heat-shock factor proteins are related to the mechanisms of SAH-associated injury, including increased inflammation and decreased neurogenesis ^72^, and vitamin B12 can be a modulator of HIF-1(Heat shock proteins) ^73^. It has been found that regulating *Rgs1, Hes1, Plp1* and *Hspa1a* could delay or attenuate the progress of inflammatory disease and SAH.^74–77^ These genes, including *Hspa1a*, were mainly found to have expression changes in SAM, IAM and PAM compared with normal microglia, indicating the possible microglia activation mechanism in these clusters. Enrichment shows these genes are involved in cell death and the immune process. Genes enrichment is shown in **Table 2**.

These results emphasize the existence of SAM, IAM and PAM under SAH conditions, which is different from normal microglial phenotype. The integrated analysis further revealed the possible targets of microglial transformation and activation after SAH. These targets and biomarkers of SAH microglia are of great biological significance in post-SAH neuroinflammation, thus helping in figuring the microglia-mediated neuroinflammation mechanism in SAH.

### 3.8. Trajectory state analysis showed the SAH microglia were all in an activated state

We applied BEAM implemented in Monocle 2 for the corresponding pseudotime analysis of normal microglia and SAH microglia (**Figure S20**). The results revealed 2 states of normal microglia, while there was only one activated state of SAH microglia.

In chronological order, the SAH microglia were IMG-C8, C4, C1, C2, C0, C6, C9, C3, C5 and C7 respectively. This suggested that SAM (IMG-C6), IAM (IMG-C3) and PAM (IMG-C7) appeared later, which indicates that the premier groups may impact the expression of these three groups. SAM may be a key subset that could induce IAM and PAM. In the transformation process, significantly increased genes including *Apoe, Actb, Atp5g2, Atp5l, Abca1, Atp6v0c, Adamts1, Actg* and *Atf3*. These genes play an important role in the activation and functional change of microglia activation from the normal state to active state.

## 4. Discussion

We report the first study to explore post-SAH microglia using scRNA-seq, revealing the diversified microglia transcriptomes post-SAH. Among microglia in the SAH-affected brain, three distinct subsets of SAM, IAM and PAM were found, whereby the molecular features of SAM and IAM differed from the previously reported activated microglial states^37, 78^. Each of the post-SAH microglia subtypes, SAM, IAM and PAM, have their unique features. SAM are characterized by high expression of *Lpl, Apoe* and *Cst7.*, while IAM exhibit upregulated *Il1a, Ccl3* and *Ccl4*. Some of these markers have been identified in previous study on AD, MS and aging disease^34, 37–39, 79^. while other marker genes of these microglial subsets appeared closely related to the progression of SAH. Of note, some genes expressed in developing microglia were found to be up-regulated in the post-SAH state, including the proliferation genes *Birc5, Mki67* and *Fabp5* in (PAM), metabolic activity genes *Mif* (SMG-C5), and *Ms4a6c* (SMG-C5) ^24, 80^. The re-expression of microglial developmental genes has also been found in previous studies on other disease models, which may represent a special way microglia adopt to cope with stress, and potentially play an important role in the activation of microglia and the pathogenesis of SAH ^24, 38^. These evidence indicates that microglial have different transcription in SAH especially on DAM, IAM and PAM. Therefore, our data not only revealed the heterogeneity of microglia under the SAH state but also enriched the microglial biological process.

In the integration analysis, it was found that the genes related to SAM (IMG-C6), IAM (IMG-C3) and PAM (IMG-C7) were highly expressed only in SAH microglia, but not in normal microglia, thus further confirming the existence of these subgroups in SAH microglia. By comparison with normal microglia belonging to the same IMG subgroup of SAH microglia, genes with strong expression changes in SAM, PAM and IAM were revealed, which mediate be the key signaling pathways driving the generation of specific post-SAH microglia clusters (**Table 2**). Different gene expressions, elevations or suppression could determine the activation destinations of microglia cells. Trajectory analysis (SMG, integrated) showed that the generation of SAM was probably earlier than that of IAM and PAM. This evidence suggests that SAM may have a more critical role in neuroinflammation after SAH. Nevertheless, future studies on the function of microglial sub-clusters are needed for the further detection of SAH.

We also analyzed the interaction of microglia subsets in the SAH state and found that microglia in SAM, IAM and PAM had the most frequent inflammatory pathway connections. The elevation of signaling pathways including CCL, GALECTIN, TGFb, APP and SPP1 signaling pathways etc. were the greatest, which could affect the development and progression of neuroinflammation after SAH ^42, 52, 56–58^.

Therefore, this study is a comprehensive, quantitative comparative study of signaling pathways in microglia mediated neuroinflammation after SAH. Some of the pathways have been mentioned in previous studies^20–22, 50, 81–86^ to modulate neuroinflammation in SAH, while we also revealed new candidate signaling pathways involved. These may serve as potential molecular targets to control post-SAH microglia activation and inflammation.

While our results emphasized the importance of microglia in neuroinflammation, other cells like CNS-associated macrophage, astrocyte, neuron, monocyte and macrophage, could also be strongly involved in neuroinflammatory responses after SAH. The interplay between brain cells should also be considered in future SAH studies. ^59^ Our summary of the interaction pathways of microglia subsets and other cell types could provide a theoretical basis for future research. For example, the function of each microglial subset and their marker genes/proteins will need to be tested experimentally by using tools such as genetic manipulation etc.

Microglial cells are an attractive target for developing biomarkers and monitoring clinical progress because they can respond sensitively to changes in the brain before physical symptoms appear ^87^. In the experimental and clinical research of SAH, it has been found that more and more microglia-related markers and regulatory factors are closely related to the prognosis of SAH ^20, 75, 88–91^. In fact, evidence of aberrant microglia activation has been found in various forms of SAH and neurodegenerative disease, but the diversity of these reactions and specificity of the microglia’s response to each disease including SAH need to be further explored ^13, 21, 90, 92, 93^. Our findings will refurbish the understanding of the pathological process of microglia mediated neuroinflammation in SAH, enhance our understandings on the physiological characteristics of microglia’s response to different diseases, and potentially accelerate the development of microglial inflammation modulators for SAH.

## 5. Supporting Information

Supporting Information is available from the Wiley Online Library or from the author.

## ACKNOWLEDGMENTS AUTHOR CONTRIBUTIONS

Junfan CHEN and Kwok Chu George WONG designed the study. Junfan CHEN carried out most of the experiments, bioinformatics analysis, manuscript writing and figure preparation. Lei SUN, Hao LYU, Huasheng LAI, Zhiyuan ZHENG performed the SAH model establishment and some of the immunostaining. Lei SUN, Yisen ZHANG and Yang WANG assisted in bioinformatics analysis. Yujie LUO, Gang LU and Wai Yee CHAN contributed to the conceptualization of the study. Xinyi CHEN and Zhongqi LI helped with the microglial isolation steps. Junfan CHEN, Ho KO and Kwok Chu George WONG wrote the manuscript with input from all authors.

We acknowledge that some of the schematics were created using BioRender (https://biorender.com/).

## ETHICS APPROVAL

The study was approved by the Animal Experimentation Ethics Committee, the Chinese University of Hong Kong (reference number 19-108-GRF).

## DATA AVAILABILITY STATEMENT

Experiment data are available upon reasonable requests to the lead authors.

## DECLARATION OF INTERESTS

The authors declare no competing interests.

## Notes

### Competing Interest Statement

The authors have declared no competing interest.

## Reference

1. Cahill J, Zhang JH. Subarachnoid hemorrhage: is it time for a new direction? Stroke. Mar 2009;40(3 Suppl):S86-7. doi:10.1161/strokeaha.108.533315

2. Johnston SC, Selvin S, Gress DR. The burden, trends, and demographics of mortality from subarachnoid hemorrhage. Neurology. May 1998;50(5):1413–8. doi:10.1212/wnl.50.5.1413

3. Wong GK, Wong R, Mok V, Wong A, Poon WS. Natural history and medical treatment of cognitive dysfunction after spontaneous subarachnoid haemorrhage: review of current literature with respect to aneurysm treatment. Journal of the neurological sciences. Dec 15 2010;299(1-2):5–8. doi:10.1016/j.jns.2010.08.059

4. Lawton MT, Vates GE. Subarachnoid Hemorrhage. The New England journal of medicine. Jul 20 2017;377(3):257–266. doi:10.1056/NEJMcp1605827

5. Wong GK, Lam S, Ngai K, Wong A, Mok V, Poon WS. Evaluation of cognitive impairment by the Montreal cognitive assessment in patients with aneurysmal subarachnoid haemorrhage: prevalence, risk factors and correlations with 3 month outcomes. Journal of neurology, neurosurgery, and psychiatry. Nov 2012;83(11):1112–7. doi:10.1136/jnnp-2012-302217

6. Wong GK, Lam SW, Ngai K, et al. Cognitive domain deficits in patients with aneurysmal subarachnoid haemorrhage at 1 year. Journal of neurology, neurosurgery, and psychiatry. Sep 2013;84(9):1054–8. doi:10.1136/jnnp-2012-304517

7. Durmaz R, Ozkara E, Kanbak G, et al. Nitric oxide level and adenosine deaminase activity in cerebrospinal fluid of patients with subarachnoid hemorrhage. Turkish neurosurgery. Apr 2008;18(2):157–64.

8. Yatsushige H, Calvert JW, Cahill J, Zhang JH. Limited role of inducible nitric oxide synthase in blood-brain barrier function after experimental subarachnoid hemorrhage. Journal of neurotrauma. Dec 2006;23(12):1874–82. doi:10.1089/neu.2006.23.1874

9. Zheng VZ, Wong GKC. Neuroinflammation responses after subarachnoid hemorrhage: A review. Journal of clinical neuroscience : official journal of the Neurosurgical Society of Australasia. Aug 2017;42:7–11. doi:10.1016/j.jocn.2017.02.001

10. van Dijk BJ, Vergouwen MD, Kelfkens MM, Rinkel GJ, Hol EM. Glial cell response after aneurysmal subarachnoid hemorrhage - Functional consequences and clinical implications. Biochimica et biophysica acta. Mar 2016;1862(3):492–505. doi:10.1016/j.bbadis.2015.10.013

11. Colonna M, Butovsky O. Microglia Function in the Central Nervous System During Health and Neurodegeneration. Annual review of immunology. Apr 26 2017;35:441–468. doi:10.1146/annurev-immunol-051116-052358

12. Prinz M, Priller J, Sisodia SS, Ransohoff RM. Heterogeneity of CNS myeloid cells and their roles in neurodegeneration. Nature neuroscience. Sep 27 2011;14(10):1227–35. doi:10.1038/nn.2923

13. Schneider UC, Davids AM, Brandenburg S, et al. Microglia inflict delayed brain injury after subarachnoid hemorrhage. Acta neuropathologica. Aug 2015;130(2):215–31. doi:10.1007/s00401-015-1440-1

14. Nayak D, Roth TL, McGavern DB. Microglia development and function. Annual review of immunology. 2014;32:367–402. doi:10.1146/annurev-immunol-032713-120240

15. Kooijman E, Nijboer CH, van Velthoven CT, Kavelaars A, Kesecioglu J, Heijnen CJ. The rodent endovascular puncture model of subarachnoid hemorrhage: mechanisms of brain damage and therapeutic strategies. Journal of neuroinflammation. Jan 3 2014;11:2. doi:10.1186/1742-2094-11-2

16. Sehba FA. Rat endovascular perforation model. Translational stroke research. Dec 2014;5(6):660–8. doi:10.1007/s12975-014-0368-4

17. Leclerc JL, Garcia JM, Diller MA, et al. A Comparison of Pathophysiology in Humans and Rodent Models of Subarachnoid Hemorrhage. Frontiers in molecular neuroscience. 2018;11:71. doi:10.3389/fnmol.2018.00071

18. Satomi J, Hadeishi H, Yoshida Y, Suzuki A, Nagahiro S. Histopathological Findings in Brains of Patients Who Died in the Acute Stage of Poor-grade Subarachnoid Hemorrhage. Neurologia medico-chirurgica. Dec 15 2016;56(12):766–770. doi:10.2176/nmc.oa.2016-0061

19. Yao X, Liu S, Ding W, et al. TLR4 signal ablation attenuated neurological deficits by regulating microglial M1/M2 phenotype after traumatic brain injury in mice. Journal of neuroimmunology. Sep 15 2017;310:38-45. doi:10.1016/j.jneuroim.2017.06.006

20. Peng J, Pang J, Huang L, et al. LRP1 activation attenuates white matter injury by modulating microglial polarization through Shc1/PI3K/Akt pathway after subarachnoid hemorrhage in rats. Redox biology. Feb 2019;21:101121. doi:10.1016/j.redox.2019.101121

21. Li R, Liu W, Yin J, et al. TSG-6 attenuates inflammation-induced brain injury via modulation of microglial polarization in SAH rats through the SOCS3/STAT3 pathway. Journal of neuroinflammation. Aug 20 2018;15(1):231. doi:10.1186/s12974-018-1279-1

22. Gris T, Laplante P, Thebault P, et al. Innate immunity activation in the early brain injury period following subarachnoid hemorrhage. Journal of neuroinflammation. Dec 4 2019;16(1):253. doi:10.1186/s12974-019-1629-7

23. Rincon F, Rossenwasser RH, Dumont A. The epidemiology of admissions of nontraumatic subarachnoid hemorrhage in the United States. Neurosurgery. Aug 2013;73(2):217–22; discussion 212-3. doi:10.1227/01.neu.0000430290.93304.33

24. Ochocka N, Segit P, Walentynowicz K, et al. Single-cell RNA sequencing reveals functional heterogeneity of glioma-associated brain macrophages. Nature communications. 2021;12(1):1151. doi:10.1038/s41467-021-21407-w

25. Okada R, Yamamoto K, Matsumoto N. DCIR3 and DCIR4 are widely expressed among tissue-resident macrophages with the exception of microglia and alveolar macrophages. Biochemistry and biophysics reports. Dec 2020;24:100840. doi:10.1016/j.bbrep.2020.100840

26. Butovsky O, Weiner H. Microglial signatures and their role in health and disease. Nature reviews Neuroscience. 2018;19(10):622–635. doi:10.1038/s41583-018-0057-5

27. Matcovitch-Natan O, Winter DR, Giladi A, et al. Microglia development follows a stepwise program to regulate brain homeostasis. Science (New York, NY). Aug 19 2016;353(6301):aad8670. doi:10.1126/science.aad8670

28. Suzuki K, Shinohara M, Uno Y, et al. Deletion of B-cell translocation gene 2 (BTG2) alters the responses of glial cells in white matter to chronic cerebral hypoperfusion. Apr 3 2021;18(1):86. doi:10.1186/s12974-021-02135-w

29. Bharti V, Tan H, Zhou H, Wang JF. Txnip mediates glucocorticoid-activated NLRP3 inflammatory signaling in mouse microglia. Neurochemistry international. Dec 2019;131:104564. doi:10.1016/j.neuint.2019.104564

30. von Rüden EL, Wolf F, Keck M, et al. Genetic Modulation of HSPA1A Accelerates Kindling Progression and Exerts Pro-convulsant Effects. Neuroscience. Aug 21 2018;386:108–120. doi:10.1016/j.neuroscience.2018.06.031

31. Zheng Z, Kim JY, Ma H, Lee JE, Yenari MA. Anti-inflammatory effects of the 70 kDa heat shock protein in experimental stroke. Journal of cerebral blood flow and metabolism : official journal of the International Society of Cerebral Blood Flow and Metabolism. Jan 2008;28(1):53–63. doi:10.1038/sj.jcbfm.9600502

32. Ekimova IV, Plaksina DV, Pastukhov YF, et al. New HSF1 inducer as a therapeutic agent in a rodent model of Parkinson’s disease. Experimental Neurology. 2018/08/01/ 2018;306:199–208. doi:https://doi.org/10.1016/j.expneurol.2018.04.012

33. Pan C, Tian M, Zhang L, et al. lncRNA Signature for Predicting Cerebral Vasospasm in Patients with SAH: Implications for Precision Neurosurgery. Molecular therapy Nucleic acids. 2020;21:983–990. doi:10.1016/j.omtn.2020.07.028

34. Masuda T, Sankowski R, Staszewski O, Prinz M. Microglia Heterogeneity in the Single-Cell Era. Cell reports. Feb 4 2020;30(5):1271–1281. doi:10.1016/j.celrep.2020.01.010

35. He X, Zhang S, Chen J, Li D. Increased LGALS3 expression independently predicts shorter overall survival in patients with the proneural subtype of glioblastoma. May 2019;8(5):2031–2040. doi:10.1002/cam4.2075

36. Lin M, Griessenauer CJ, Starke RM, et al. Haplotype analysis of SERPINE1 gene: Risk for aneurysmal subarachnoid hemorrhage and clinical outcomes. Molecular genetics & genomic medicine. Aug 2019;7(8):e737. doi:10.1002/mgg3.737

37. Keren-Shaul H, Spinrad A, Weiner A, et al. A Unique Microglia Type Associated with Restricting Development of Alzheimer’s Disease. Cell. Jun 15 2017;169(7):1276–1290.e17. doi:10.1016/j.cell.2017.05.018

38. Hammond TR, Dufort C, Dissing-Olesen L, et al. Single-Cell RNA Sequencing of Microglia throughout the Mouse Lifespan and in the Injured Brain Reveals Complex Cell-State Changes. Immunity. Jan 15 2019;50(1):253–271.e6. doi:10.1016/j.immuni.2018.11.004

39. Li Q, Cheng Z, Zhou L, et al. Developmental Heterogeneity of Microglia and Brain Myeloid Cells Revealed by Deep Single-Cell RNA Sequencing. Neuron. Jan 16 2019;101(2):207–223.e10. doi:10.1016/j.neuron.2018.12.006

40. Hagemeyer N, Hanft KM, Akriditou MA, et al. Microglia contribute to normal myelinogenesis and to oligodendrocyte progenitor maintenance during adulthood. Acta neuropathologica. Sep 2017;134(3):441–458. doi:10.1007/s00401-017-1747-1

41. Wlodarczyk A, Holtman I, Krueger M, et al. A novel microglial subset plays a key role in myelinogenesis in developing brain. The EMBO journal. 2017;36(22):3292–3308. doi:10.15252/embj.201696056

42. Lanterna LA, Rigoldi M, Tredici G, et al. APOE influences vasospasm and cognition of noncomatose patients with subarachnoid hemorrhage. Neurology. 2005;64(7):1238–1244. doi:10.1212/01.wnl.0000156523.77347.b4

43. Krasemann S, Madore C, Cialic R, et al. The TREM2-APOE Pathway Drives the Transcriptional Phenotype of Dysfunctional Microglia in Neurodegenerative Diseases. Immunity. Sep 19 2017;47(3):566–581.e9. doi:10.1016/j.immuni.2017.08.008

44. Calandra T, Roger T. Macrophage migration inhibitory factor: a regulator of innate immunity. Nature reviews Immunology. Oct 2003;3(10):791–800. doi:10.1038/nri1200

45. Kannan-Thulasiraman P, Seachrist DD, Mahabeleshwar GH, Jain MK, Noy N. Fatty acid-binding protein 5 and PPARbeta/delta are critical mediators of epidermal growth factor receptor-induced carcinoma cell growth. The Journal of biological chemistry. Jun 18 2010;285(25):19106–15. doi:10.1074/jbc.M109.099770

46. Liu RZ, Mita R, Beaulieu M, Gao Z, Godbout R. Fatty acid binding proteins in brain development and disease. The International journal of developmental biology. 2010;54(8-9):1229–39. doi:10.1387/ijdb.092976rl

47. Bruce KD, Gorkhali S, Given K, et al. Lipoprotein Lipase Is a Feature of Alternatively-Activated Microglia and May Facilitate Lipid Uptake in the CNS During Demyelination. Frontiers in molecular neuroscience. 2018;11:57–57. doi:10.3389/fnmol.2018.00057

48. Gao Y, Vidal-Itriago A, Kalsbeek MJ, et al. Lipoprotein Lipase Maintains Microglial Innate Immunity in Obesity. Cell reports. Sep 26 2017;20(13):3034–3042. doi:10.1016/j.celrep.2017.09.008

49. Jordão M, Sankowski R, Brendecke S, et al. Single-cell profiling identifies myeloid cell subsets with distinct fates during neuroinflammation. Science (New York, NY). 2019;363(6425)doi:10.1126/science.aat7554

50. Kroner A, Greenhalgh AD, Zarruk JG, Passos Dos Santos R, Gaestel M, David S. TNF and increased intracellular iron alter macrophage polarization to a detrimental M1 phenotype in the injured spinal cord. Neuron. Sep 3 2014;83(5):1098-116. doi:10.1016/j.neuron.2014.07.027

51. Li S, Yang S, Sun B, Hang C. Melatonin attenuates early brain injury after subarachnoid hemorrhage by the JAK-STAT signaling pathway. International journal of clinical and experimental pathology. 2019;12(3):909–915.

52. Nishikawa H, Nakatsuka Y, Shiba M, Kawakita F, Fujimoto M, Suzuki H. Increased Plasma Galectin-3 Preceding the Development of Delayed Cerebral Infarction and Eventual Poor Outcome in Non-Severe Aneurysmal Subarachnoid Hemorrhage. Translational stroke research. Apr 2018;9(2):110–119. doi:10.1007/s12975-017-0564-0

53. Füger P, Hefendehl JK, Veeraraghavalu K, et al. Microglia turnover with aging and in an Alzheimer’s model via long-term in vivo single-cell imaging. Nature neuroscience. 2017/10/01 2017;20(10):1371–1376. doi:10.1038/nn.4631

54. Mathys H, Adaikkan C, Gao F, et al. Temporal Tracking of Microglia Activation in Neurodegeneration at Single-Cell Resolution. Cell reports. 2017;21(2):366–380. doi:10.1016/j.celrep.2017.09.039

55. O’Neil SM, Witcher KG, McKim DB, Godbout JP. Forced turnover of aged microglia induces an intermediate phenotype but does not rebalance CNS environmental cues driving priming to immune challenge. Nov 26 2018;6(1):129. doi:10.1186/s40478-018-0636-8

56. Tada T, Kanaji M, Kobayashi S. Induction of communicating hydrocephalus in mice by intrathecal injection of human recombinant transforming growth factor-beta 1. Journal of neuroimmunology. Mar 1994;50(2):153–8. doi:10.1016/0165-5728(94)90041-8

57. Nakatsuka Y, Kawakita F, Yasuda R, et al. Preventive effects of cilostazol against the development of shunt-dependent hydrocephalus after subarachnoid hemorrhage. Journal of neurosurgery. Aug 2017;127(2):319–326. doi:10.3171/2016.5.jns152907

58. Nishikawa H, Liu L, Nakano F, et al. Modified Citrus Pectin Prevents Blood-Brain Barrier Disruption in Mouse Subarachnoid Hemorrhage by Inhibiting Galectin-3. Stroke. Nov 2018;49(11):2743–2751. doi:10.1161/strokeaha.118.021757

59. Prinz M, Jung S, Priller J. Microglia Biology: One Century of Evolving Concepts. Cell. Oct 3 2019;179(2):292–311. doi:10.1016/j.cell.2019.08.053

60. Liu Y, Zhang H, Li Y, et al. Long Noncoding RNA Rps4l Mediates the Proliferation of Hypoxic Pulmonary Artery Smooth Muscle Cells. Hypertension (Dallas, Tex : 1979). 2020;76(4):1124–1133. doi:10.1161/hypertensionaha.120.14644

61. Zhao H, Li S, Zhu J, Hua X, Wan L. Analysis of Peripheral Blood Cells’ Transcriptome in Patients With Subarachnoid Hemorrhage From Ruptured Aneurysm Reveals Potential Biomarkers. World neurosurgery. 2019;129:e16–e22. doi:10.1016/j.wneu.2019.04.125

62. Crotti A, Ransohoff R. Microglial Physiology and Pathophysiology: Insights from Genome-wide Transcriptional Profiling. Immunity. 2016;44(3):505–515. doi:10.1016/j.immuni.2016.02.013

63. Vikman P, Beg S, Khurana T, Hansen-Schwartz J, Edvinsson L. Gene expression and molecular changes in cerebral arteries following subarachnoid hemorrhage in the rat. Journal of Neurosurgery JNS. 01 Sep. 2006 2006;105(3):438-444. doi:10.3171/jns.2006.105.3.438

64. Chen J, Jin J, Zhang X, et al. Microglial lnc-U90926 facilitates neutrophil infiltration in ischemic stroke via MDH2/CXCL2 axis. Molecular Therapy. 2021/09/01/ 2021;29(9):2873–2885. doi:https://doi.org/10.1016/j.ymthe.2021.04.025

65. Subedi L, Lee JH, Yumnam S, Ji E, Kim SY. Anti-Inflammatory Effect of Sulforaphane on LPS-Activated Microglia Potentially through JNK/AP-1/NF-κB Inhibition and Nrf2/HO-1 Activation. Cells. 2019;8(2):194.

66. Dunn LL, Kong SMY, Tumanov S, et al. Hmox1 (Heme Oxygenase-1) Protects Against Ischemia-Mediated Injury via Stabilization of HIF-1α (Hypoxia-Inducible Factor-1α). Arteriosclerosis, thrombosis, and vascular biology. Jan 2021;41(1):317–330. doi:10.1161/atvbaha.120.315393

67. Schallner N, Pandit R, LeBlanc R, et al. Microglia regulate blood clearance in subarachnoid hemorrhage by heme oxygenase-1. The Journal of clinical investigation. 2015;125(7):2609–25. doi:10.1172/jci78443

68. Monif M, Reid CA, Powell KL, Drummond KJ, O’Brien TJ, Williams DA. Interleukin-1β has trophic effects in microglia and its release is mediated by P2X7R pore. Journal of neuroinflammation. 2016/06/30 2016;13(1):173. doi:10.1186/s12974-016-0621-8

69. Olah M, Menon V, Habib N, et al. Single cell RNA sequencing of human microglia uncovers a subset associated with Alzheimer’s disease. Nature communications. 2020;11(1):6129. doi:10.1038/s41467-020-19737-2

70. Grosche L, Knippertz I, König C, et al. The CD83 Molecule – An Important Immune Checkpoint. Review. Frontiers in Immunology. 2020-April-17 2020;11(721)doi:10.3389/fimmu.2020.00721

71. Jeon H, Kim JH, Kim JH, Lee WH, Lee MS, Suk K. Plasminogen activator inhibitor type 1 regulates microglial motility and phagocytic activity. Journal of neuroinflammation. Jun 29 2012;9:149. doi:10.1186/1742-2094-9-149

72. Zuo Y, Wang J, Liao F, et al. Inhibition of Heat Shock Protein 90 by 17-AAG Reduces Inflammation via P2X7 Receptor/NLRP3 Inflammasome Pathway and Increases Neurogenesis After Subarachnoid Hemorrhage in Mice. Frontiers in molecular neuroscience. 2018;11:401. doi:10.3389/fnmol.2018.00401

73. Ghemrawi R, Pooya S, Lorentz S, et al. Decreased vitamin B12 availability induces ER stress through impaired SIRT1-deacetylation of HSF1. Cell death & disease. 2013;4:e553. doi:10.1038/cddis.2013.69

74. Zuo Y, Wang J, Liao F, et al. Inhibition of Heat Shock Protein 90 by 17-AAG Reduces Inflammation via P2X7 Receptor/NLRP3 Inflammasome Pathway and Increases Neurogenesis After Subarachnoid Hemorrhage in Mice. Frontiers in molecular neuroscience. 2018;11

75. Lee JK, Bou Dagher J. Regulator of G-protein Signaling (RGS)1 and RGS10 Proteins as Potential Drug Targets for Neuroinflammatory and Neurodegenerative Diseases. The AAPS journal. May 2016;18(3):545–9. doi:10.1208/s12248-016-9883-4

76. Groh J, Klein D, Berve K, West B, Martini R. Targeting microglia attenuates neuroinflammation-related neural damage in mice carrying human PLP1 mutations. Glia. 2019;67(2):277–290. doi:10.1002/glia.23539

77. Zou W, Chen Q-X, Sun X-W, et al. Acupuncture inhibits Notch1 and Hes1 protein expression in the basal ganglia of rats with cerebral hemorrhage. Neural Regen Res. 2015;10(3):457–462. doi:10.4103/1673-5374.153696

78. Saijo K, Glass CK. Microglial cell origin and phenotypes in health and disease. Nature reviews Immunology. Oct 25 2011;11(11):775–87. doi:10.1038/nri3086

79. Masuda T, Sankowski R, Staszewski O, et al. Spatial and temporal heterogeneity of mouse and human microglia at single-cell resolution. Nature. Feb 2019;566(7744):388–392. doi:10.1038/s41586-019-0924-x

80. Huang Y, Xu Z, Xiong S, et al. Repopulated microglia are solely derived from the proliferation of residual microglia after acute depletion. Nature neuroscience. 2018;21(4):530–540. doi:10.1038/s41593-018-0090-8

81. Buchanan MM, Hutchinson M, Watkins LR, Yin H. Toll-like receptor 4 in CNS pathologies. Journal of neurochemistry. Jul 2010;114(1):13–27. doi:10.1111/j.1471-4159.2010.06736.x

82. Samraj AK, Müller AH, Grell AS, Edvinsson L. Role of unphosphorylated transcription factor STAT3 in late cerebral ischemia after subarachnoid hemorrhage. Journal of cerebral blood flow and metabolism : official journal of the International Society of Cerebral Blood Flow and Metabolism. May 2014;34(5):759–63. doi:10.1038/jcbfm.2014.15

83. Venkatraman A, Hardas S, Patel N, Singh Bajaj N, Arora G, Arora P. Galectin-3: an emerging biomarker in stroke and cerebrovascular diseases. European journal of neurology. Feb 2018;25(2):238–246. doi:10.1111/ene.13496

84. Yip PK, Carrillo-Jimenez A, King P, et al. Galectin-3 released in response to traumatic brain injury acts as an alarmin orchestrating brain immune response and promoting neurodegeneration. Scientific reports. Jan 27 2017;7:41689. doi:10.1038/srep41689

85. Jeon SB, Yoon HJ, Chang CY, Koh HS, Jeon SH, Park EJ. Galectin-3 exerts cytokine-like regulatory actions through the JAK-STAT pathway. Journal of immunology (Baltimore, Md : 1950). Dec 1 2010;185(11):7037-46. doi:10.4049/jimmunol.1000154

86. Aoki T, Koseki H, Miyata H, et al. RNA sequencing analysis revealed the induction of CCL3 expression in human intracranial aneurysms. Scientific reports. 2019;9(1):10387. doi:10.1038/s41598-019-46886-2

87. Hong S, Beja-Glasser V, Nfonoyim B, et al. Complement and microglia mediate early synapse loss in Alzheimer mouse models. Science (New York, NY). 2016;352(6286):712–716. doi:10.1126/science.aad8373

88. Suzuki H, Fujimoto M, Kawakita F, et al. Toll-Like Receptor 4 and Tenascin-C Signaling in Cerebral Vasospasm and Brain Injuries After Subarachnoid Hemorrhage. Acta neurochirurgica Supplement. 2020;127:91–96. doi:10.1007/978-3-030-04615-6_15

89. Chen J, Wong GKC. Microglia accumulation and activation after subarachnoid hemorrhage. Neural Regen Res. Aug 2021;16(8):1531–1532. doi:10.4103/1673-5374.303028

90. Zheng ZV, Lyu H, Lam SYE, Lam PK, Poon WS, Wong GKC. The Dynamics of Microglial Polarization Reveal the Resident Neuroinflammatory Responses After Subarachnoid Hemorrhage. Translational stroke research. Jun 2020;11(3):433-449. doi:10.1007/s12975-019-00728-5

91. Gris T, Laplante P, Thebault P, et al. Innate immunity activation in the early brain injury period following subarachnoid hemorrhage. Journal of neuroinflammation. 2019;16(1):253. doi:10.1186/s12974-019-1629-7

92. Lan X, Han X, Li Q, Yang QW, Wang J. Modulators of microglial activation and polarization after intracerebral haemorrhage. Nature reviews Neurology. Jul 2017;13(7):420–433. doi:10.1038/nrneurol.2017.69

93. Salter MW, Stevens B. Microglia emerge as central players in brain disease. Nature Medicine. 2017/09/01 2017;23(9):1018–1027. doi:10.1038/nm.4397

