## Supplementary material for "Single-cell analysis of microglial transcriptomic diversity in subarachnoid hemorrhage": Supporting information- V40.docx

Kwok Chu George WONG*

**Table S1: Key resources**

| **Reagent** | **Manufacturer** | **Cat. number** |
| --- | --- | --- |
| Anti- CD16/32 | BD Biosciences | 553141 |
| Anti- CD206 | R&D,MMR | AF2535 |
| Tmem119 Monoclonal Antibody, Alexa Fluor 488 | Invitrogen | 53-6119-80 |
| ART 1000G genomic racked ster | Thermo Fisher Scientifi | 2019G |
| Bovine Serum Albumin | Sigma | A9418-50G |
| CD11b-APC | Miltenyi Biotec | 130-113-793 |
| CD11b MicroBeads, mouse | Miltenyi Biotec | 130-049-601 |
| Cell strainers 70um | Falcon | 352350 |
| DNase I, grade II | Roche | 10104159001 |
| Gentle MACS 25 C Tubes | Miltenyi Biotec | 130-090-237 |
| Goat serum | Abcam | ab7481 |
| IBA1 | Abcam | Ab5076 |
| LS Separation columns | Miltenyi Biotec | 130-042-401 |
| Neural Tissue Dissociation Kit- Papain | Miltenyi Biotec | 130-092-628 |
| PBS | Thermo Fisher Scientifi | 10010031 |
| Propidium Iodide Solution | Miltenyi Biotec | 130-093-233 |
| RNASin(R) plus RNAse inhibitor | Promega-gm | N2615 |
| RNAseZAP | Thermo Fisher Scientific | AM9780 |
| Trypan Blue | Abcam | Ab233465 |
| **Equipment** | **Manufacturer** | **Cat. number** |
| QuadroMACS Separator | Miltenyi Biotec | 130-090-976 |
| MACS Multistand | Miltenyi Biotec | 130-042-303 |
| gentleMACS Dissociator | Miltenyi Biotec | 130-093-235 |
| BD LSRFortessa™ Cell Analyzer | BD Biosciences | |
| Nikon Eclipse Ti Inverted Microscope | Nikon | |

**I: Supplemental Methods:**

1. **Animals**:

C57BL/6 (wild type) were obtained from the Laboratory Animal Services Centre of the Chinese University of Hong Kong (male, 12 weeks, average weight 25–30g). The mouse was placed in separate ventilation cages and exposed to food and water freely at 23^o^C and 50-60% humidity in a 12-hour/12-hour day and night cycle. All procedures involving animals and their care have been approved by the Ethics Committee of the Chinese University of Hong Kong.

2. **SAH model perforate:**

Prematurity subarachnoid hemorrhage (SAH) model was adopted in this study. ^1^^,^^2^ In simple: mice were anesthetized and fixed in a supine position. A surgical microscope was used in the whole process of model establishment. Firstly, a 1 cm incision was made in the midline of the mice neck, and then the left common carotid artery (CCA), left external carotid artery (ECA) and left internal carotid artery (ICA) were dissected clearly. ECA was ligated at the distal end, and two 1.5-cm-length 5–0 silk sutures were prepared for filament fixation. Block the blood of ECA and insert the filament (20-mm-long blunted 5–0 monofilament nylon suture) to ECA and ICA continue to the intracranial vessels. The vessel was perforated at the bifurcation of the middle cerebral artery (MCA) where the resistance was encountered. Then, the filament was immediately pulled out and then the hemorrhage was introduced into subarachnoid space. The sham model with the same procedure except for filament perforation. During the whole operation and recovery process, the mice were kept at 37 °C. To protect their eyesight, their eyes were coated with ointment. Buprenorphine was given intraperitoneally (i.p.) for analgesia twice a day for 3 days. On the day 1st, 3rd, 5th, and 10^th^ day after SAH induction, the body weight of mice was evaluated for wellbeing.

3. **Phenotype evaluation:**

Motor capacities was evaluated on the 1, 3, 5 and 10 days after SAH (nSHAM = nSAH = 6-8). To confirm SAH induction using two phenotypic tests: the holding time test and the Modified Bederson Score. The evaluating investigator was blinded to the experimental conditions. The holding time test is adapted from the inverted grid test and has been widely used in SAH model assessment.^3^^,^^4^ Briefly, a cotton-tipped applicator was placed and fixed on a pedestal at a 30 ° angle. Then, the mice were placed on it, and the time for the mice to remain in suspension was measured. Each mouse was measured three times to get the average time. The Modified Bederson score ^5^^,^^6^ was applied to evaluate neurological function. The mouse model of Moderate SAH model (Modified Bederson Score 2-3) with the holding time test at D1 (range of 23.33±10.69) was recruited for the microglia study.

**Table S2: Modified Bederson Score**

| **Modified Bederson Score** | **Description** |
| --- | --- |
| 4 | Longitudinal spinning or seizure activity. |
| 3 | Unidirectional circling |
| 2 | As for 1, plus decreased resistance to lateral push. |
| 1 | Forelimb flexion. |
| 0 | No deficit. |

4. **Immunohistochemistry:**

Immunohistochemistry (IHC) was used to examine the condition of microglia. The paraffin brain sections (5 µm) were firstly through a xylene/ethanol dewax-rehydration series, then antigen retrieval was performed with citrate buffer for 20 minutes. Then, after the incubation of the first antibody and second antibody, endogenous peroxidize activity was quenched with 0.3% Hydrogen peroxide (H_2_O_2_). The brain slices were prepared in blocking buffer containing 2.5% goat serum, 1% Bovine serum albumin (BSA) for one hour and the primary antibody Iba1 (1:200; Abcam, #ab5076) was applied subsequently at 4 °C overnight. Envision+System-Horseradish peroxidase (HRP) secondary antibody was applied for 1 hour at room temperature. Finally, Diaminobenzidine (DAB) was utilized. Six random fields were examined on Cortex adjacent to the perforated site (CAPS), Hippocampus (HIP) (The CA1 region of the hippocampus was selected for analysis), and Motor cortex (M1 cortex) (Left and right) respectively of each mouse under Microscope (Nikon) at 20X magnification. Microglial cell count was quantified by Image-Pro software.

5. **Immunofluorescence:**

Immunofluorescence (IF) was performed to define the post-SAH microglial polarization. Frozen sections were used for the IF, in simple, Mice were cardinally perfused with PBS followed by 10% buffered formalin then dehydration with the gradient concentration of sucrose solution from 15% to 30% and then embedded in the Optimal cutting temperature (OCT) compound for cryosection. The frozen sections were immunolabeled with primary antibodies including CD16/32 (1:200; BD Biosciences, #553141), and CD206 (1:500; R&D, MMR, #AF2535), at 4 °C overnight. Fluorescence-conjugated secondary antibodies were then incubated with frozen sections, including Donkey anti - Rat Donkey DyLight 680 IgG H+L (1:200; Invitrogen, #SA5-10030), anti-Goat Alexa Fluor® 647 IgG H+L (1:200; Invitrogen, #A21447), and Donkey anti-Rat Alexa Fluor® 488 IgG H+L (1:200; Invitrogen, #A21208), at room temperature for 2 hours. Then washed with PBS and mounted with 4′,6-diamidino-2-phenylindole (DAPI) (Abcam, #ab104139). Immunofluorescent images were acquired using a microscope (Nikon Eclipse Ti Inverted Microscope, Nikon). Quantification of M1/M2 microglial phenotype was carried out in three randomly selected high power microscopic fields across three sections.

6. **Acute microglial isolation and purification:**

Centrifuges and tools are prechilled to 4℃ or on ice. Mice (3-5 mice usually, for this study was four) at 3^rd^-day post-SAH was anesthetized and then transcardially perfused with cold PBS for 2–4 minutes each mice using a 30 ml syringe with a 20 Gauge needle. Then quickly dissect the brains, put them in cold PBS, and wash them twice to remove the blood, hair and fiber. The olfactory bulb and the cerebellum were removed. Neural Tissue Dissociation Kit P was used for brain digestion. Cut brain into small pieces with sterile scissors, and put it into a gentleMACS™ C tube containing prewarmed enzyme solution for mechanical dissociation. Then, the C tube was inverted on the gentleMACS™ Dissociator, the program (Mouse brain program) was run, and reagents were added according to the commercial protocol (Miltenyibiotec) and published paper. ^7-9^ Pass cell digest over a 70 μm filter (Pre-wetted with 1ml PBS/BSA 0.5%) to remove cell clumps, then set into a 50 ml conical tube. Spin at 400 × g(RCF) and 4℃ for 10 minutes and aspirate supernatant. Added adequate CD11b (Microglia) MicroBeads based on the cell number and incubated 15 minutes at 4℃. LS column and QuadroMACS were used for the positive selection of microglia. (1 LS column: Max. number of labeled cells: 2×10⁷) After positive selection, take out the LS column from the magnetic holder then push the solution through the LS column with the plunger provided. This will apply gentle pressure to remove the microglia from the LS column and to obtain Microglia Fraction. Centrifuge the Microglia Fraction at 300g for 10 minutes at 4°C and discard the supernatant. Put cell in 1.5ml EP tube also with 10ml 0.5% BSA-PBS buffer with 2μL RNAse inhabitor at 4 ^0^C. ^7-10^

7. **Flow Cytometry:**

Microglia suspension was washed with 0.5% BSA-PBS and followed with cell surface staining at 4 °C for 30min using the following markers: Tmem119 Monoclonal Antibody-Alexa Fluor 488 (Invitrogen, #53-6119-80), CD11b-APC (Miltenyi Biotec, #130-113-793), Propidium Iodide Solution (Miltenyi Biotec, #130-093-233). Then cell will be analyzed on BD LSRFortessa™ Cell Analyzer (BD Biosciences) according to the manufacturer's instructions. Cell viability was assessed using the Trypan blue (Abcam, #Ab233465) cell analysis on a hemocytometer.

8. **Single cell RNA sequencing:**

CD11b positive cell suspension purified from MACS were sequenced by the Chromium single-cell gene expression platform (10x Genomics). According to the manufacturer's instructions, about 10,000-15,000 microglia of each sample were directly loaded into each sample and then combined into droplets with barcoded beads by using the Chromium controller and Chromium Single-Cell 3′ Reagent Kits v3 (10x Genomics). According to the manufacturer's specifications, the barcode library was generated, and then the samples were sequenced to an average depth of 40,000-60,000 reads on a DNBSEQ-PE100 (BGI).

9. **Single-cell data analysis and Bioinformatics:**

Sequencing results were demultiplexed and converted to FASTQ format using Illumina bcl2fastq software. Sample demultiplexing, barcode processing and single-cell 3’ gene counting by using the Cell Ranger pipeline(https://support.10xgenomics.com/single-cell-geneexpression/ software/pipelines/latest/what-is-cell-ranger, version 3.1.0) and scRNA-seq data were aligned to Ensembl genome GRCm38 reference genome, a total of 13,194 single cell captured from 4 SAH mouse brain were processed using 10X Genomics Chromium Single Cell 3’ Solution. The Cell Ranger output was loaded into Seurat (version 3.1.1) to be used for Dimensional reduction, clustering, and analysis of scRNA-seq data. Overall, 8,916 cells passed the quality control threshold: all genes expressed in less than 1 cell were removed, the number of genes expressed per cell was > 500 as low and <4000 as high cut- off, UMI counts less than 500, the percent of mitochondrial-DNA derived gene-expression <10%. By using Tmem119 and Cx3r1 as confirmed microglia markers ^7^^,^^11^^,^^12^, 5824 microglia were involved in bioinformatics. Further, we recruit microglia transcriptome from normal adult mouse brain ScRNA-seq dataset (Microglia isolated by CD11b magnetic beads, and sequenced by 10X genomes platform (Same with our isolation and sequencing method), 4 normal male mice samples, age 14 weeks, C57BL/6, whole brain) and make integration analysis of SAH microglia (microglia number: 5854) and normal microglia (microglia number: 8160) with the same method with SAH microglia bioinformatics. For data from different experiments, Seurat CCA integration functions were used. ^7^^,^^13^ (Normal sample: GSM3442026, GSM3442027, GSM3442030 and GSM3442031)

To visualize the data, we further reduced the dimensionality of all 8,916 cells (For integration analysis were 14014 cells) using Seurat and used Uniform Manifold Approximation and Projection (UMAP) to project the cells into 2D space, The steps includes:1. Using the LogNormalize method of the "Normalization" function of the Seurat software to calculate the expression value of genes; 2. PCA (Principal component analysis) analysis was performed using the normalized expression value, Within all the PCs, the top 10 PCs were used to do clustering and UMAP analysis；3. To find clusters, select the weighted Shared Nearest Neighbor (SNN) graph-based clustering method. Marker genes for each cluster were identified with the "bimod"（Likelihood-ratio test）with default parameters via the FindAllMarkers function in Seurat. This selects markers genes which are expressed in more than 10% of the cells in a cluster and average log2 (fold change) of greater than 0.26. 4. Monocle2 was applied for the trajectory state analyzed and Cellchat was applied to analyze cell interaction. 5. Integration analysis with the same method above.

10. **Statistics:**

All the data were expressed as mean ± SEM. Statistical analyses were conducted by IBM SPSS 22.0 software. For the cross-sectional evaluation cohort, One-way ANOVA was used for statistical analysis to evaluate the microglia change across 4 time points. The independent t-test was used to determine the significance comparison between groups. The equality of error variance was tested as appropriate. P<0.05 after Bonferroni adjustment for multiple comparisons was considered statistically significant.

**Reference**

1.	Zheng ZV, Lyu H, Lam SYE, Lam PK, Poon WS, Wong GKC. The Dynamics of Microglial Polarization Reveal the Resident Neuroinflammatory Responses After Subarachnoid Hemorrhage. *Translational stroke research*. Jun 2020;11(3):433-449. doi:10.1007/s12975-019-00728-5

2.	Du G, Lu G, Zheng Z, Poon W, Wong K. Endovascular Perforation Murine Model of Subarachnoid Hemorrhage. *Acta neurochirurgica Supplement*. 2016;121:83-8. doi:10.1007/978-3-319-18497-5_14

3.	Aartsma-Rus A, van Putten M. Assessing functional performance in the mdx mouse model. *Journal of visualized experiments : JoVE*. 2014;(85)doi:10.3791/51303

4.	Gris T, Laplante P, Thebault P, et al. Innate immunity activation in the early brain injury period following subarachnoid hemorrhage. *Journal of neuroinflammation*. 2019;16(1):253. doi:10.1186/s12974-019-1629-7

5.	Ruan J, Yao Y. Behavioral tests in rodent models of stroke. *Brain hemorrhages*. 2020;1(4):171-184. doi:10.1016/j.hest.2020.09.001

6.	Lyu C, Zhang Y, Gu M, et al. IRAK-M Deficiency Exacerbates Ischemic Neurovascular Injuries in Experimental Stroke Mice. *Frontiers in cellular neuroscience*. 2018;12:504. doi:10.3389/fncel.2018.00504

7.	Hammond TR, Dufort C, Dissing-Olesen L, et al. Single-Cell RNA Sequencing of Microglia throughout the Mouse Lifespan and in the Injured Brain Reveals Complex Cell-State Changes. *Immunity*. Jan 15 2019;50(1):253-271.e6. doi:10.1016/j.immuni.2018.11.004

8.	Holt L, Stoyanof S, Olsen M. Magnetic Cell Sorting for In Vivo and In Vitro Astrocyte, Neuron, and Microglia Analysis. *Current protocols in neuroscience*. 2019;88(1):e71. doi:10.1002/cpns.71

9.	Hickman S, El Khoury J. Analysis of the Microglial Sensome. *Methods in molecular biology (Clifton, NJ)*. 2019;2034:305-323. doi:10.1007/978-1-4939-9658-2_23

10.	Sharma K, Schmitt S, Bergner C, et al. Cell type- and brain region-resolved mouse brain proteome. *Nature neuroscience*. 2015;18(12):1819-31. doi:10.1038/nn.4160

11.	Masuda T, Sankowski R, Staszewski O, Prinz M. Microglia Heterogeneity in the Single-Cell Era. *Cell reports*. Feb 4 2020;30(5):1271-1281. doi:10.1016/j.celrep.2020.01.010

12.	Li Q, Cheng Z, Zhou L, et al. Developmental Heterogeneity of Microglia and Brain Myeloid Cells Revealed by Deep Single-Cell RNA Sequencing. *Neuron*. Jan 16 2019;101(2):207-223.e10. doi:10.1016/j.neuron.2018.12.006

13.	Satija R, Farrell JA, Gennert D, Schier AF, Regev A. Spatial reconstruction of single-cell gene expression data. *Nature biotechnology*. 2015/05/01 2015;33(5):495-502. doi:10.1038/nbt.3192

**II: Supplemental Figures (Totally 20 figures)**

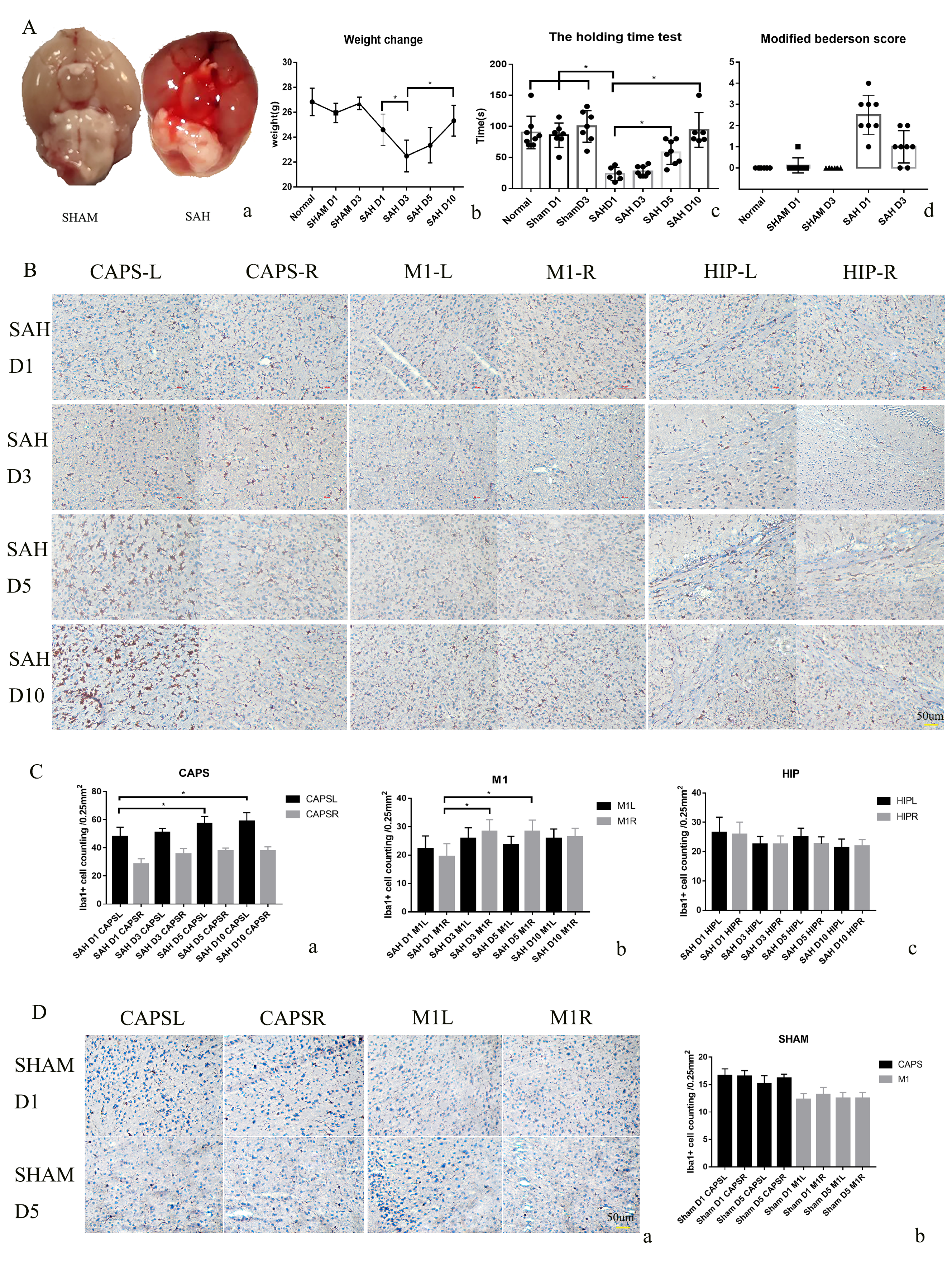

**Figure S1-** A: SAH model evaluation; B: Iba1+ IHC staining in D1, D3, D5 and D10 of SAH. (CAPS, M1 and Hippocampus); C: Quantitative analysis of Iba1+ cell; D: Iba1+ IHC staining in D1 and D5 in sham group. n=6-8, *p<0.05

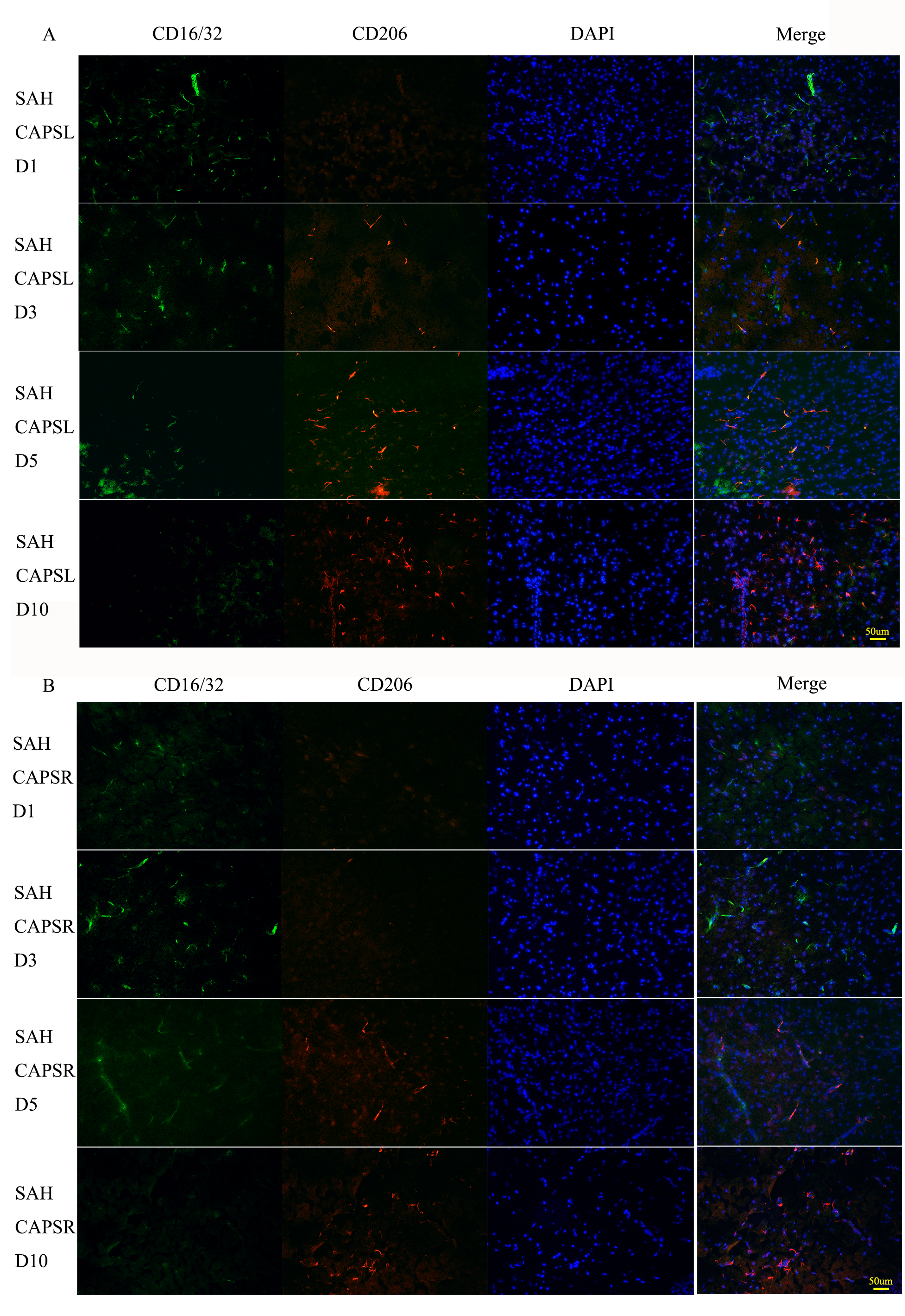

**Figure S2-** IF staining of CD16/32 (Green) and CD206 (Red), in D1, D3, D5 and D10 of SAH group in CAPS left (A) and CAPS right (B).

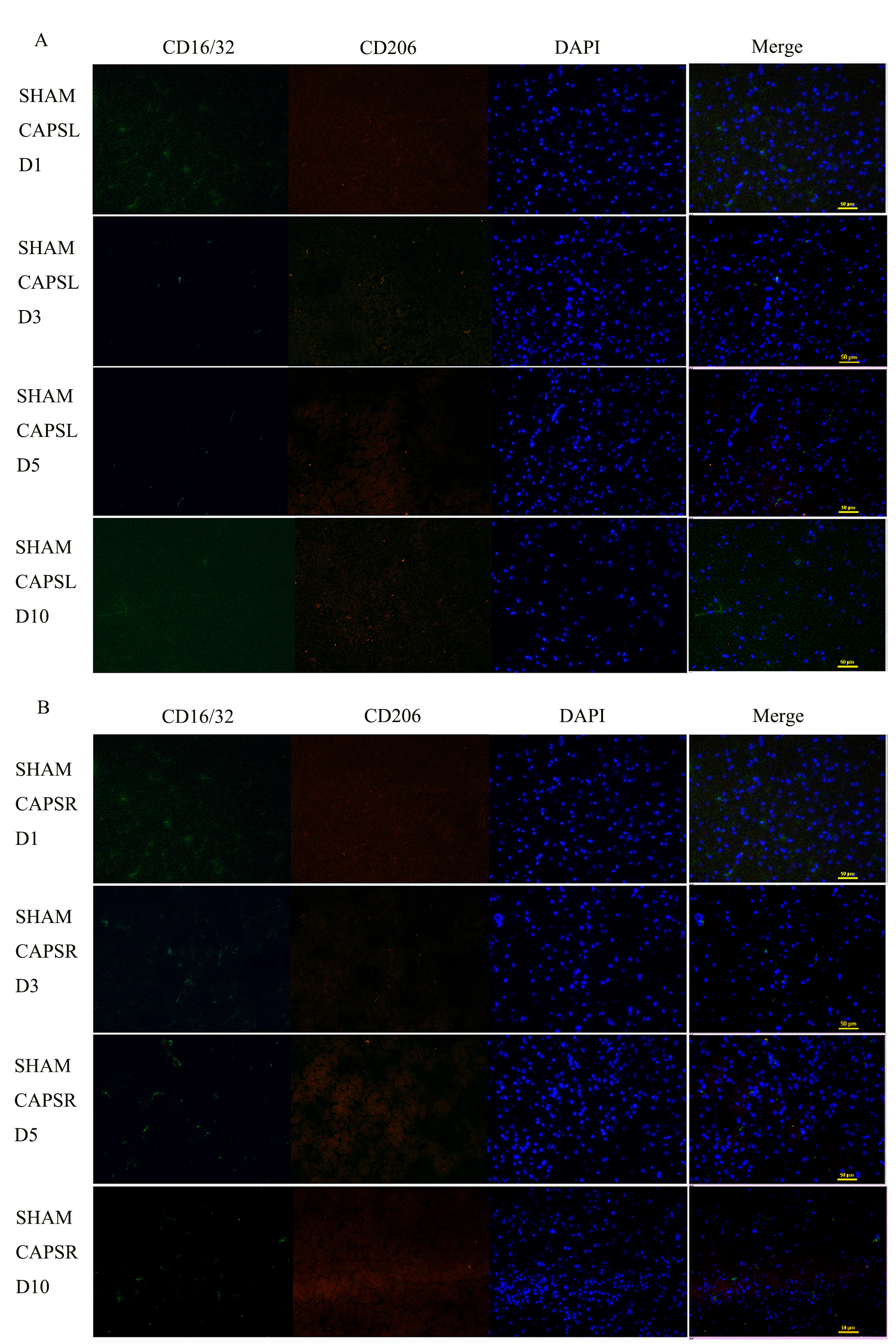

**Figure S3-** IF staining of CD16/32 (Green) and CD206 (Red), in D1, D3, D5 and D10 of SAHM group in CAPS left (A) and CAPS right (B).

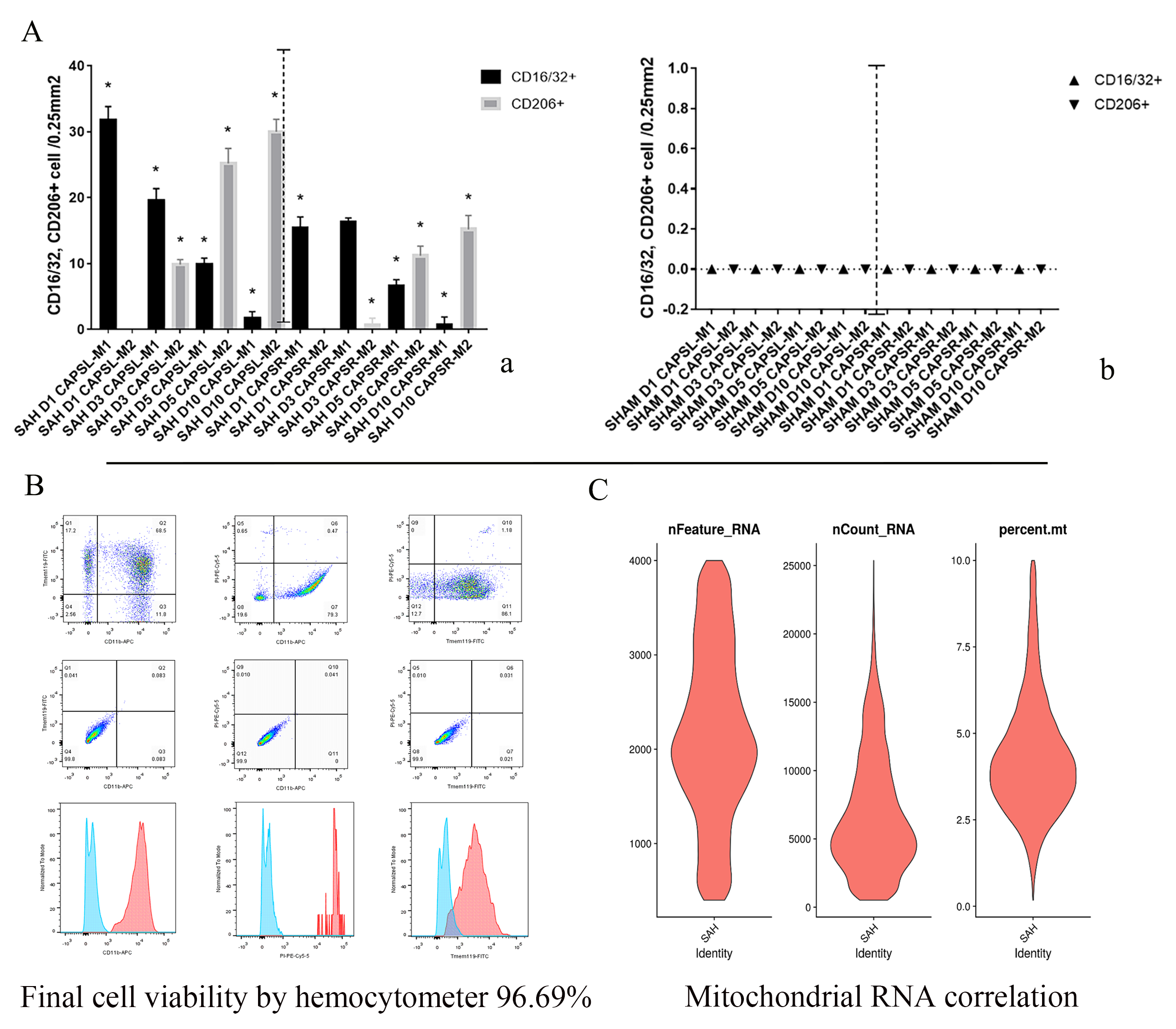

**Figure S4-** A-B: Quantitative analysis of CD16/32 and CD206 in SAH (A) and sham group (B). n=6-8, *p<0.05. C: Flow cytometry results of isolated cells; D: Mitochondrial RNA correlation results.

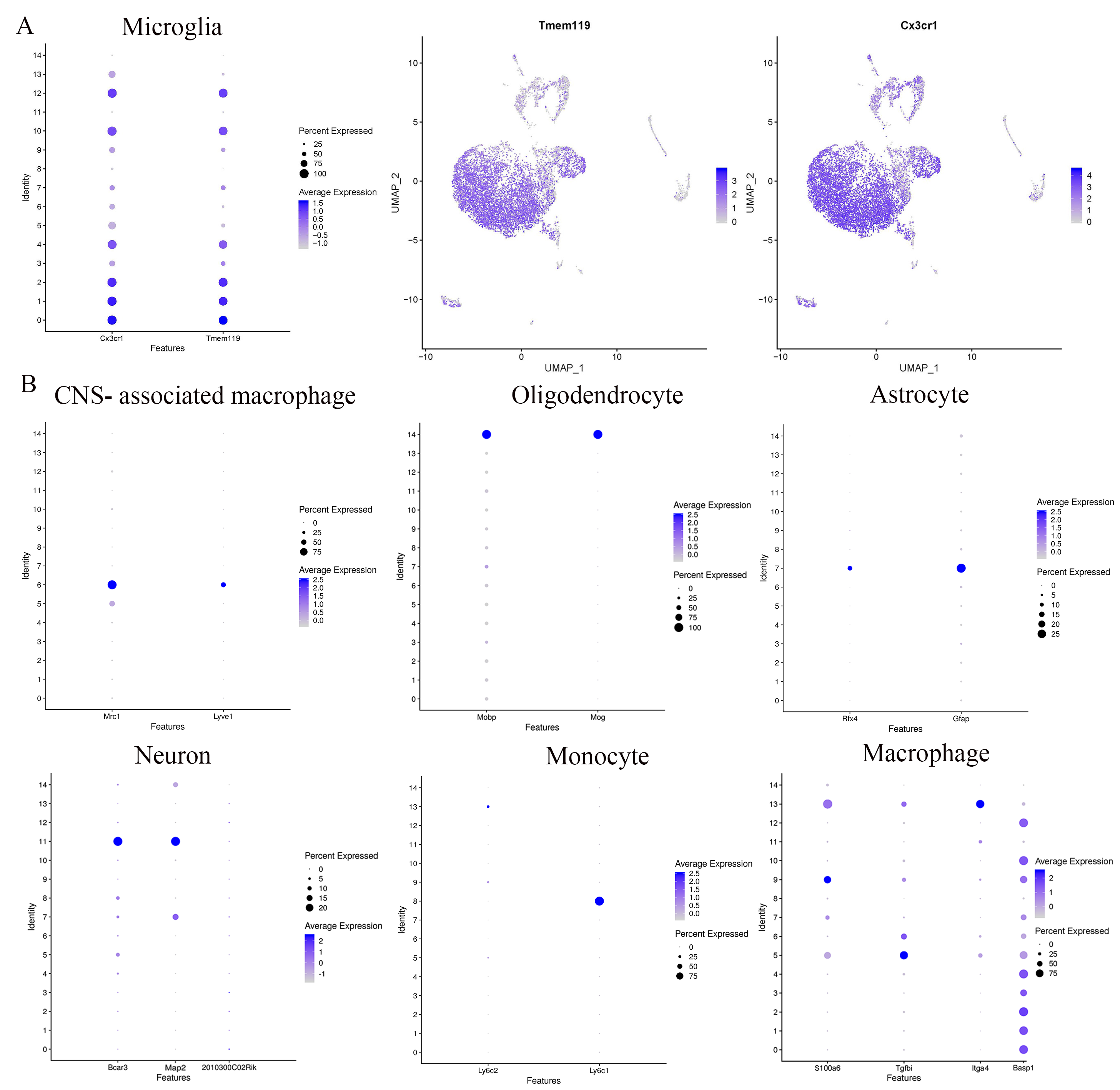

**Figure S5-** A: Microglial cell markers distribution; B: Cell markers distribution of CNS- associated macrophage, oligodendrocyte, astrocyte, neuron, monocyte and macrophage.

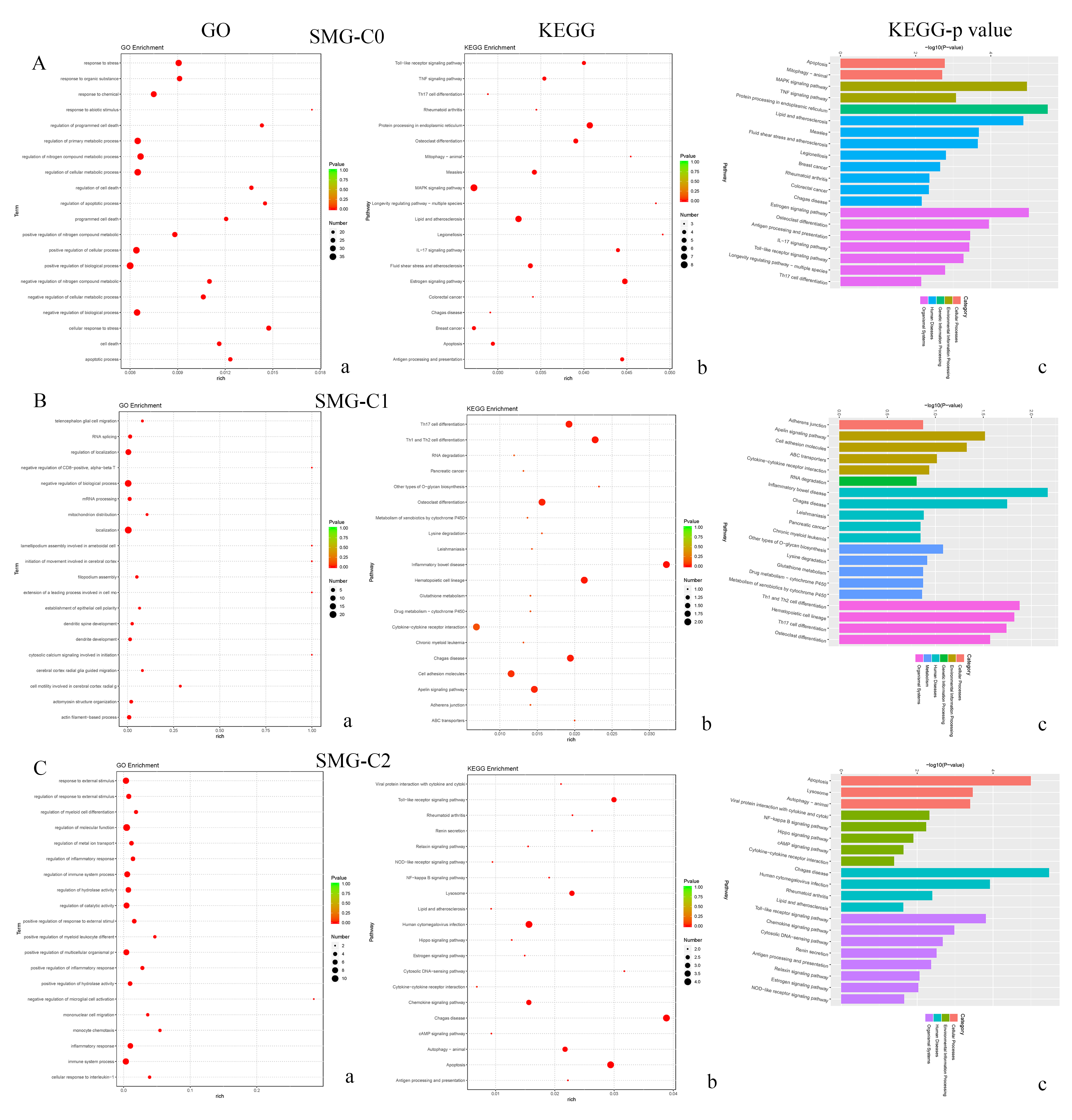

**Figure S6-** Enrichment analysis for SMG microglia. A: SMG-C0, B: SMG-C1 and C: SMG-C2; a: GO and b-c: KEGG.

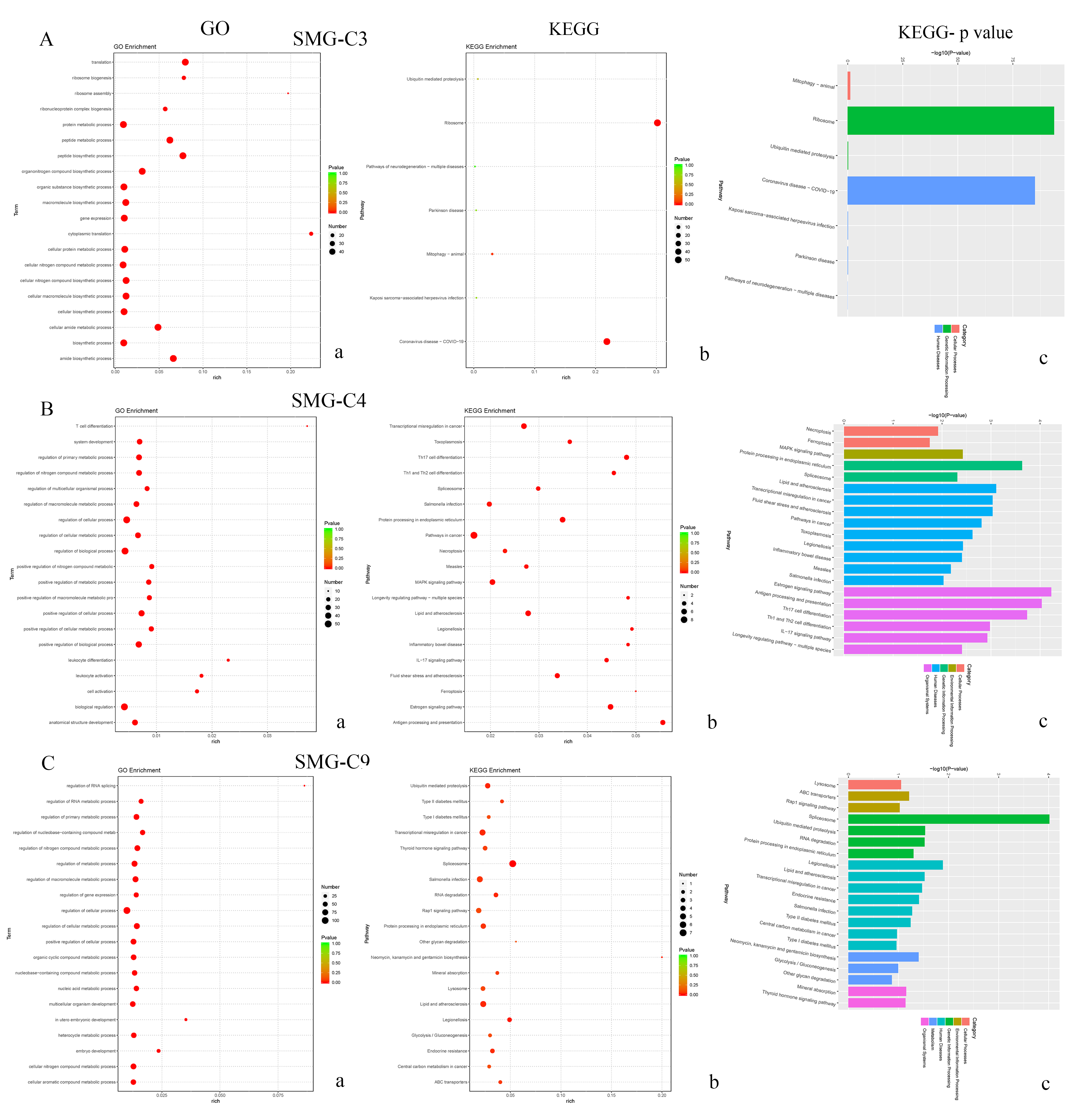

**Figure S7-** Enrichment analysis for SMG microglia. A: SMG-C3, B: SMG-C4 (B) and C: SMG-C9. a: GO and b-c: KEGG.

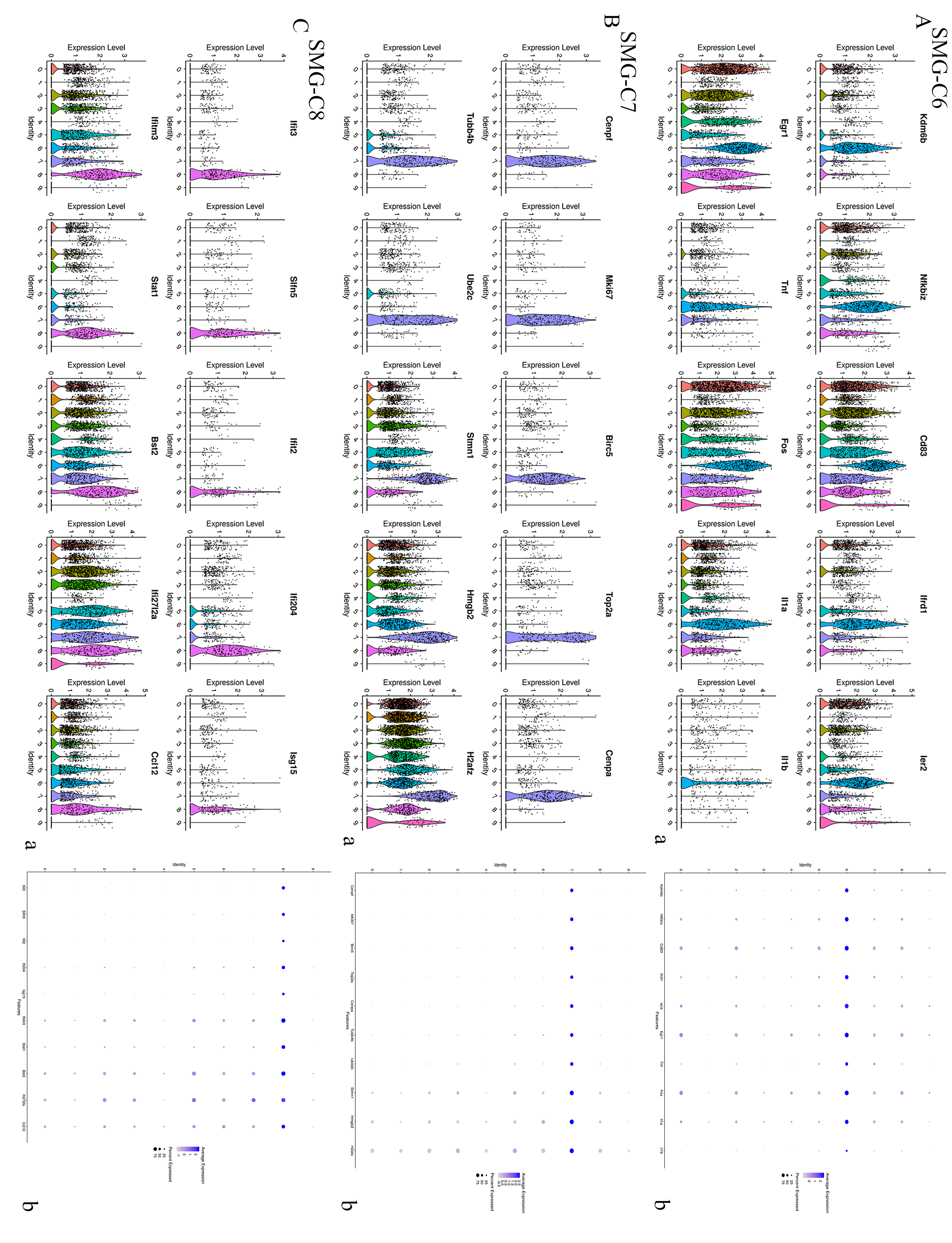

**Figure S8-** Top 10 expressed genes in SMG microglia- Atlas of violins and dot plot.

A: SMG-C6, B: SMG-C7 and C: SMG-C8.

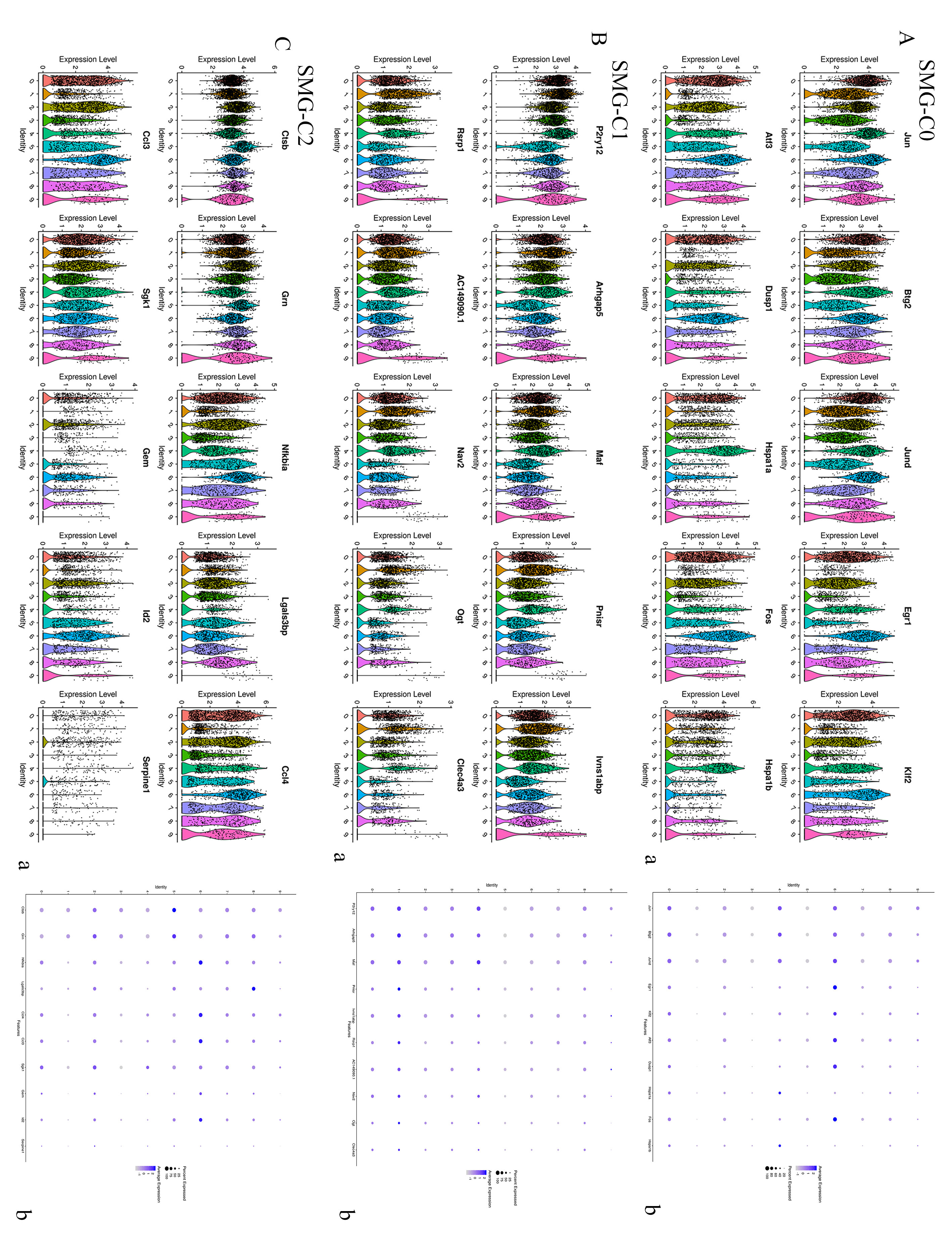

**Figure S9-** Top 10 expressed genes in SMG microglia- Atlas of violins and dot plot.

A: SMG-C0, B: SMG-C1 and C: SMG-C2.

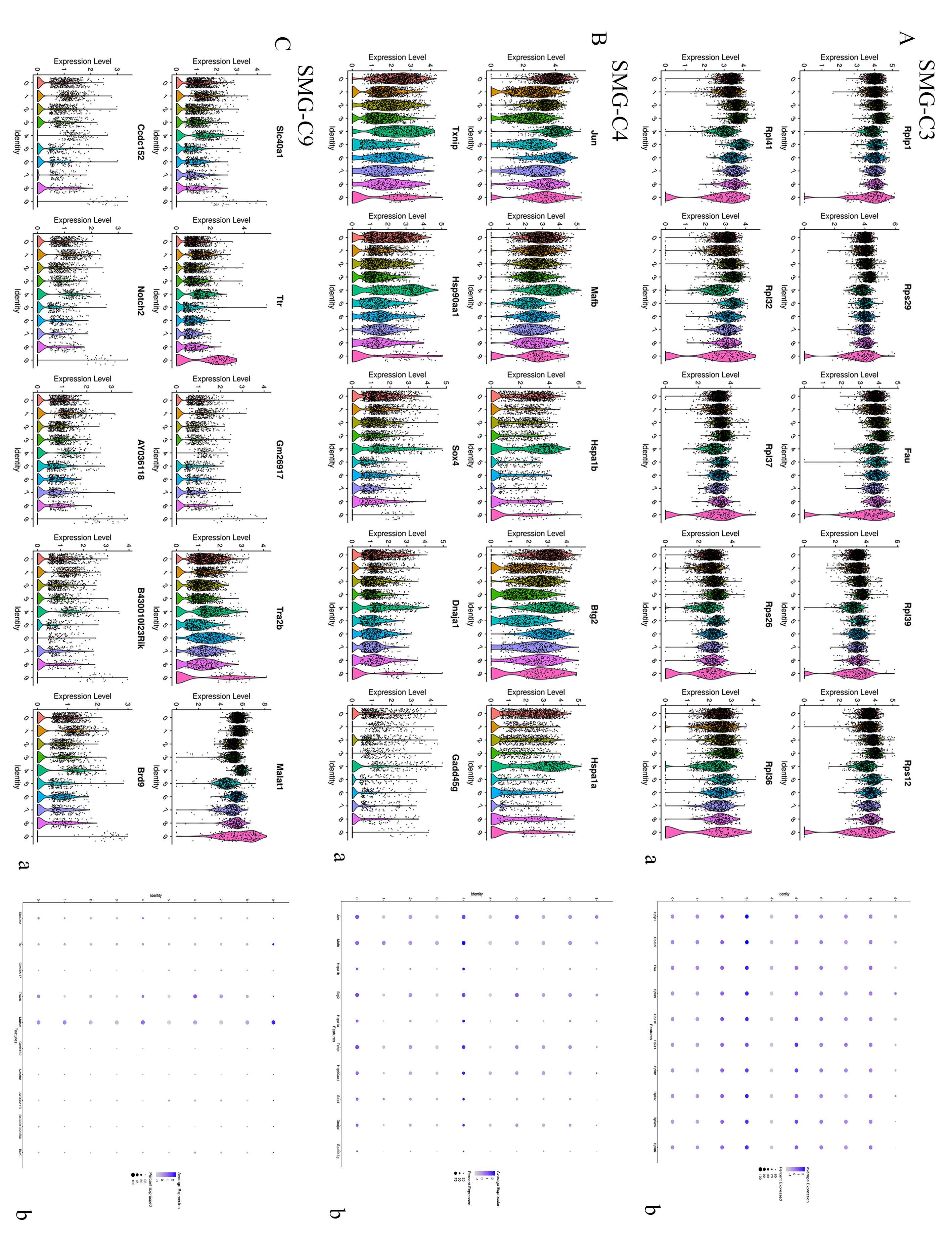

**Figure S10-** Top 10 expressed genes in SMG microglia- Atlas of violins and dot plot.

A: SMG-C3, B: SMG-C4 and C: SMG-C9.

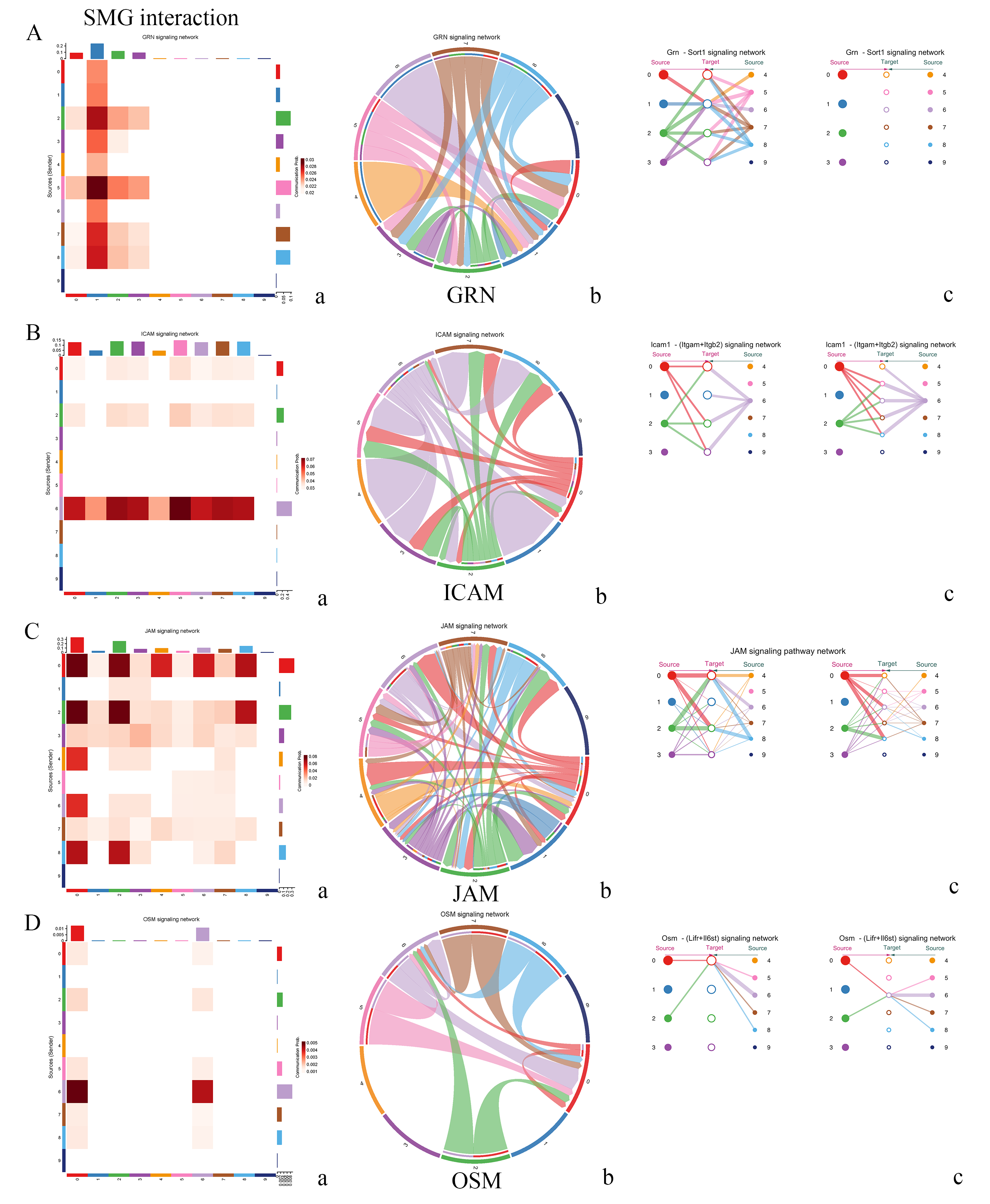

**Figure S11-** SMG microglia subsets interaction (signaling pathway). A: GRN, B: ICAM, C: JAM and D: OSM signaling pathway. a: heatmap, b: chord map and c: hierarchy connection.

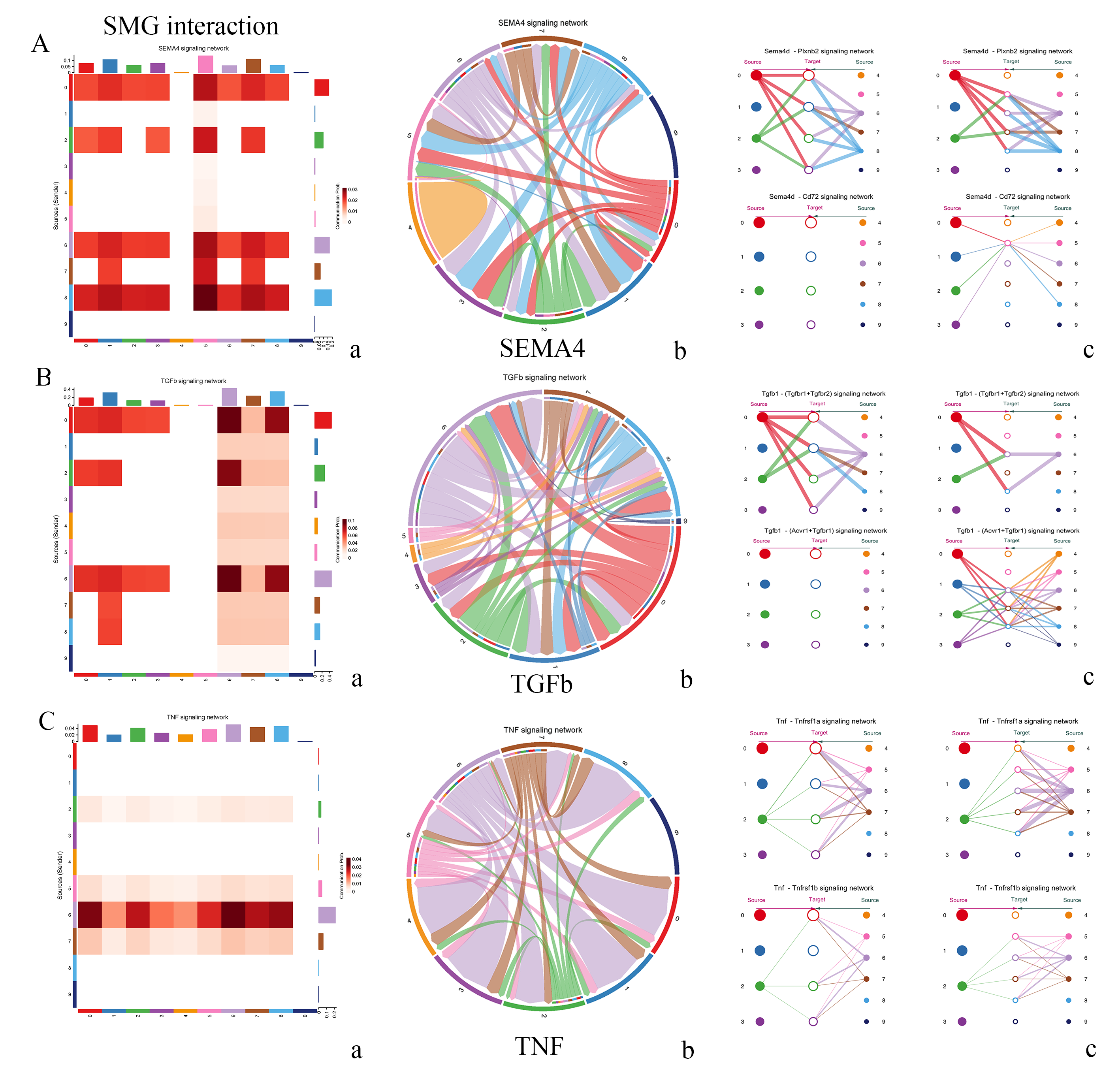

**Figure S12-** SMG microglia subsets interaction (signaling pathway). A: SEMA4, B: TGFb and C: TNF signaling pathway. a: heatmap, b; chord map and c: hierarchy connection.

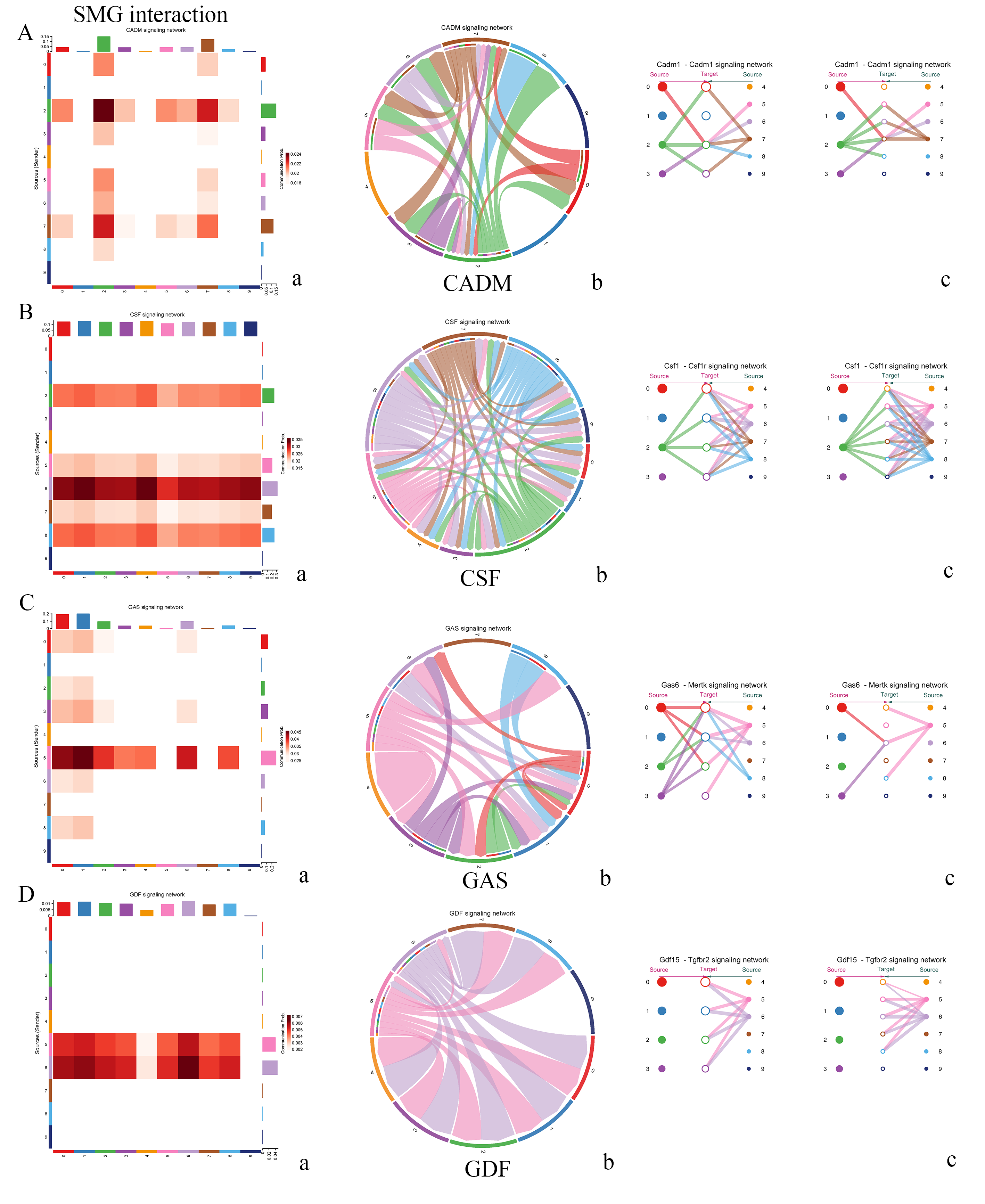

**Figure S13-** SMG microglia subsets interaction (signaling pathway) A: CADM, B: CSF, C: GAS and D: GDF signaling pathway. a: heatmap, b; chord map and c: hierarchy connection.

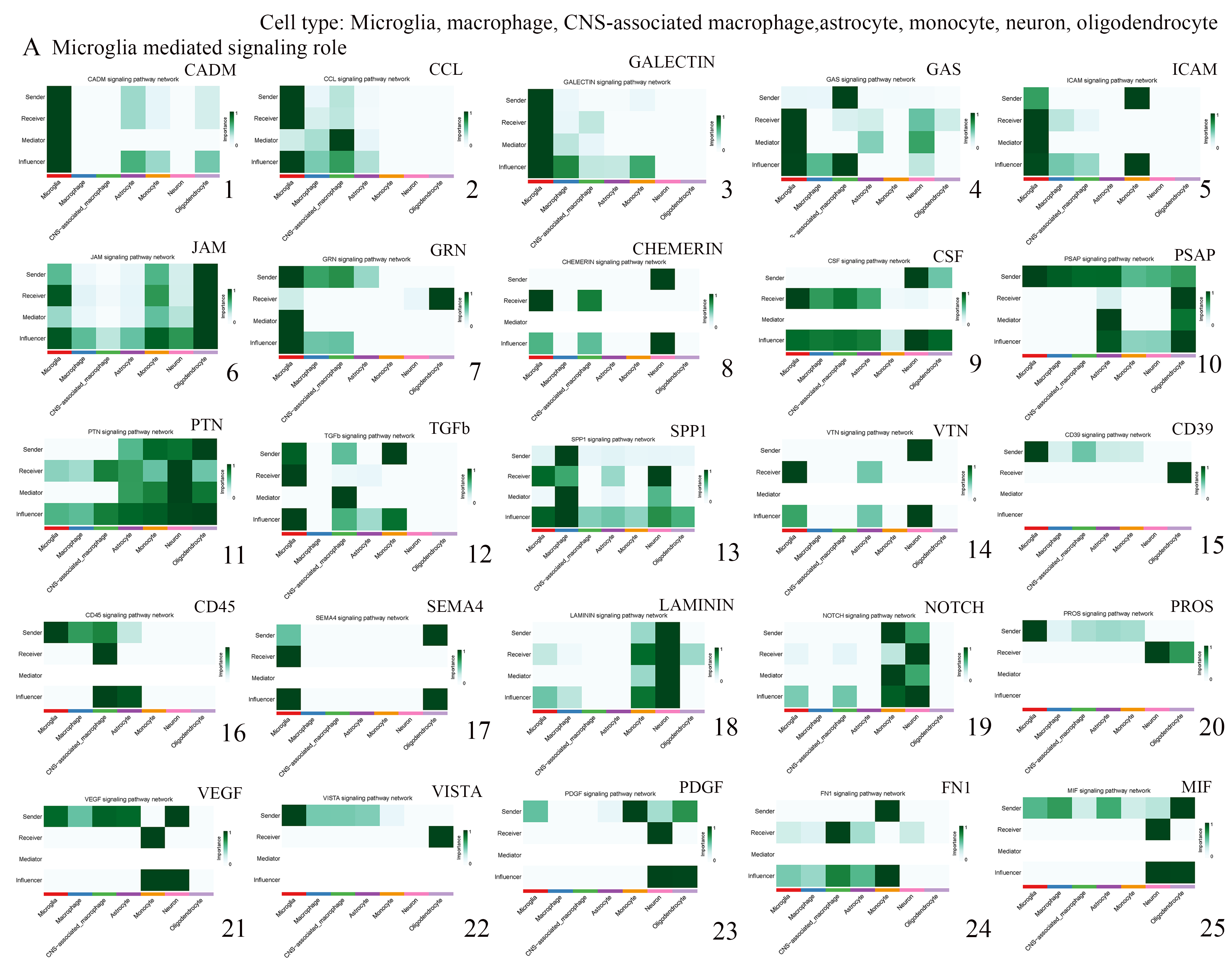

**Figure S14-** Microglia interacts with Microglia, macrophage, CNS-associated macrophage, astrocyte, monocyte, neuron and oligodendrocyte, Major contributing signaling role. A: microglia participated signaling pathway.

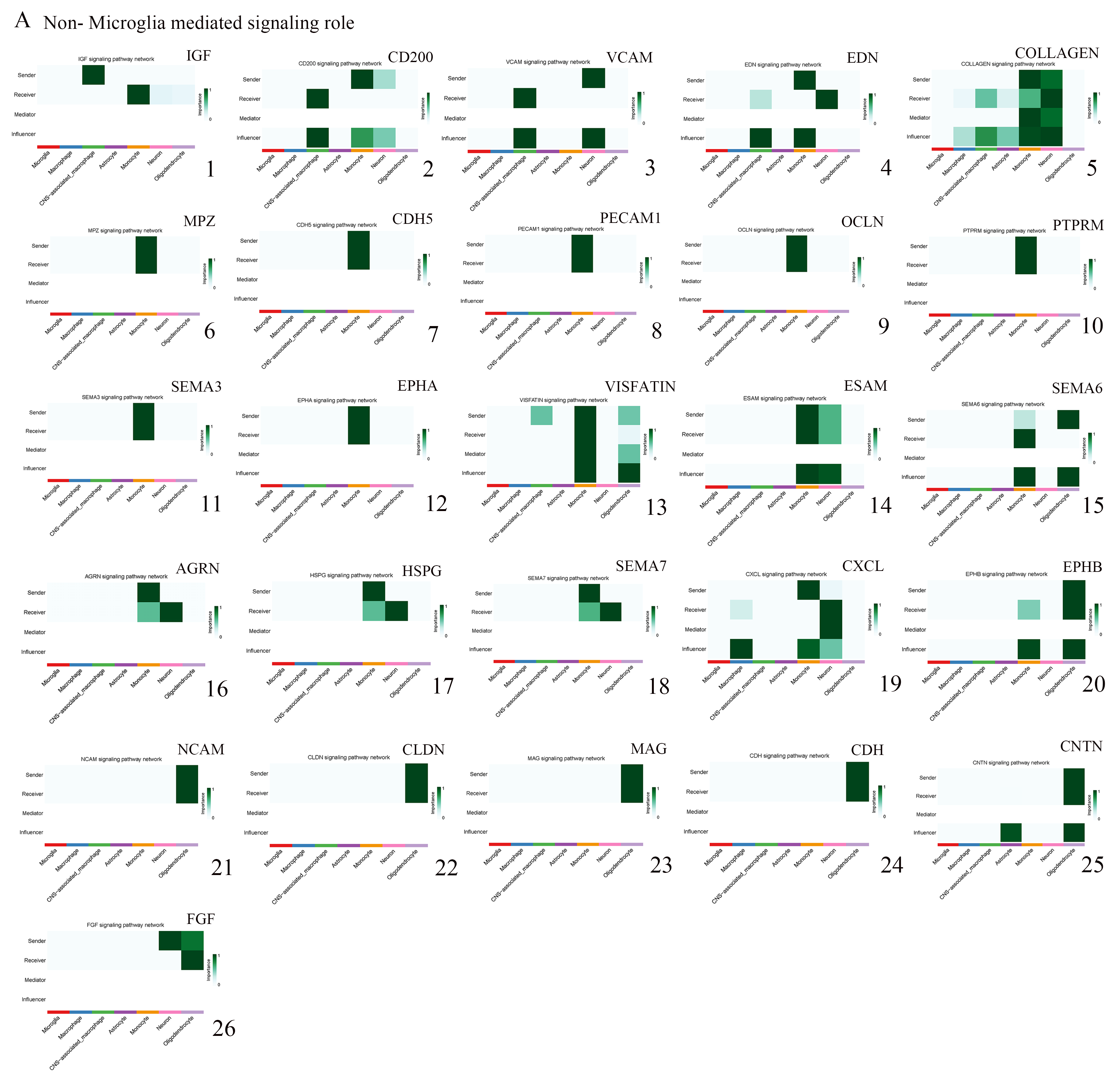

**Figure S15-** Microglia interacts with Microglia, macrophage, CNS-associated macrophage, astrocyte, monocyte, neuron and oligodendrocyte- Major contributing signaling role. A: None-microglia participated signaling pathway.

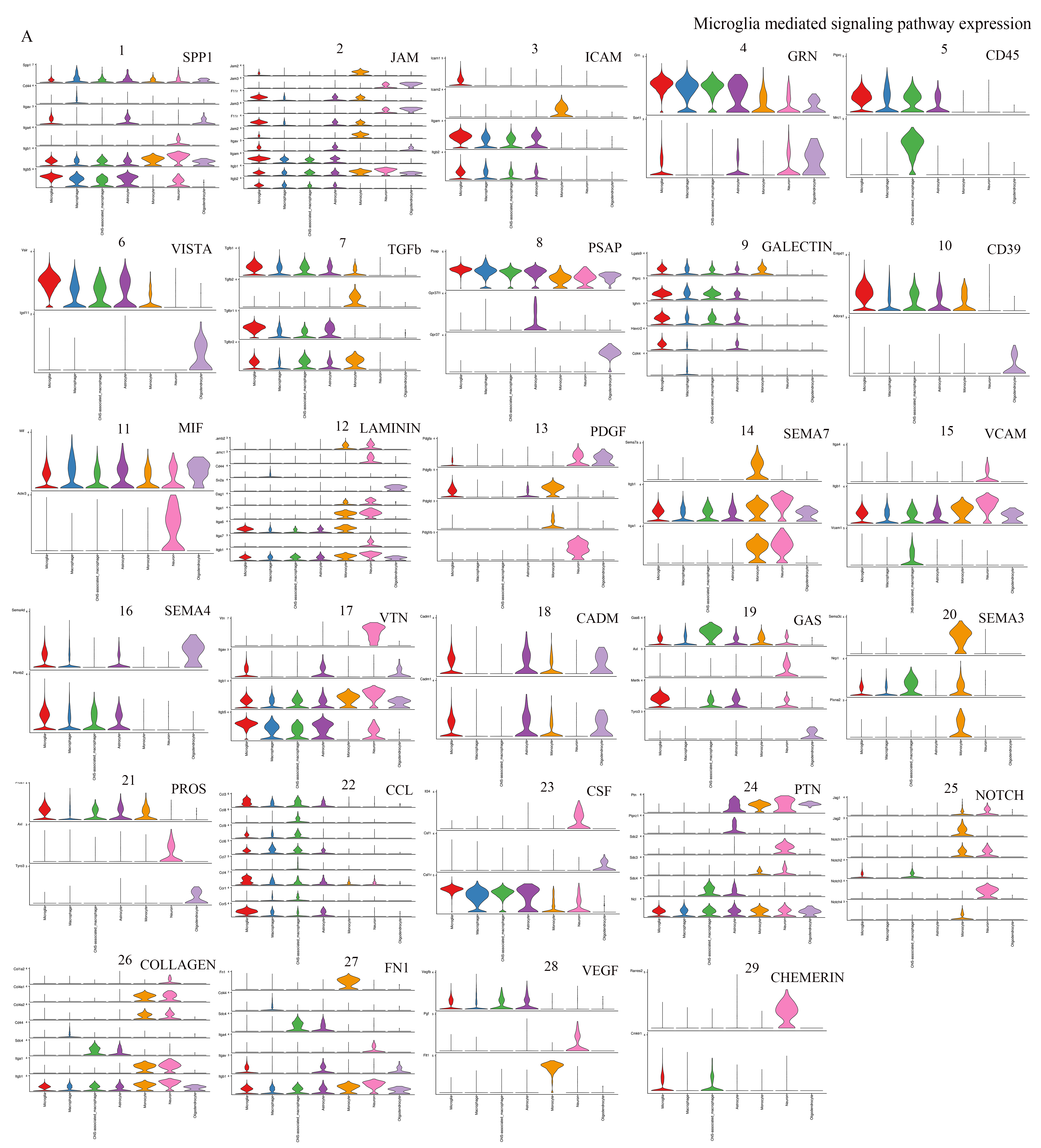

**Figure S16-** Microglia interacts with Microglia, macrophage, CNS-associated macrophage, astrocyte, monocyte, neuron and oligodendrocyte- Expression plot of signaling pathway. A: microglia participated signaling pathway.

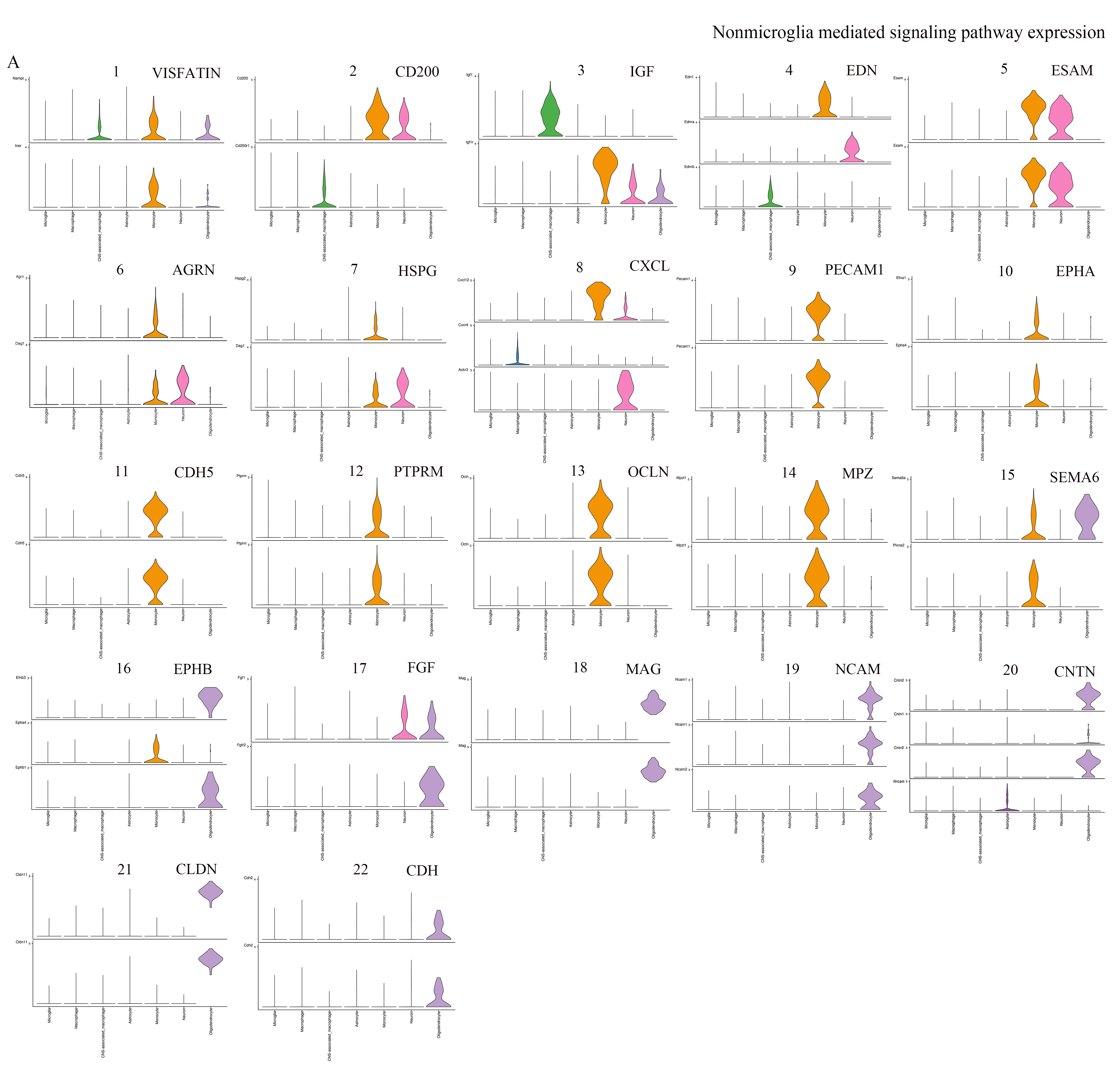

**Figure S17-** Microglia interaction with Microglia, macrophage, CNS-associated macrophage, astrocyte, monocyte, neuron and oligodendrocyte- Expression plot of signaling pathway. A: Non-microglia participated signaling pathway.

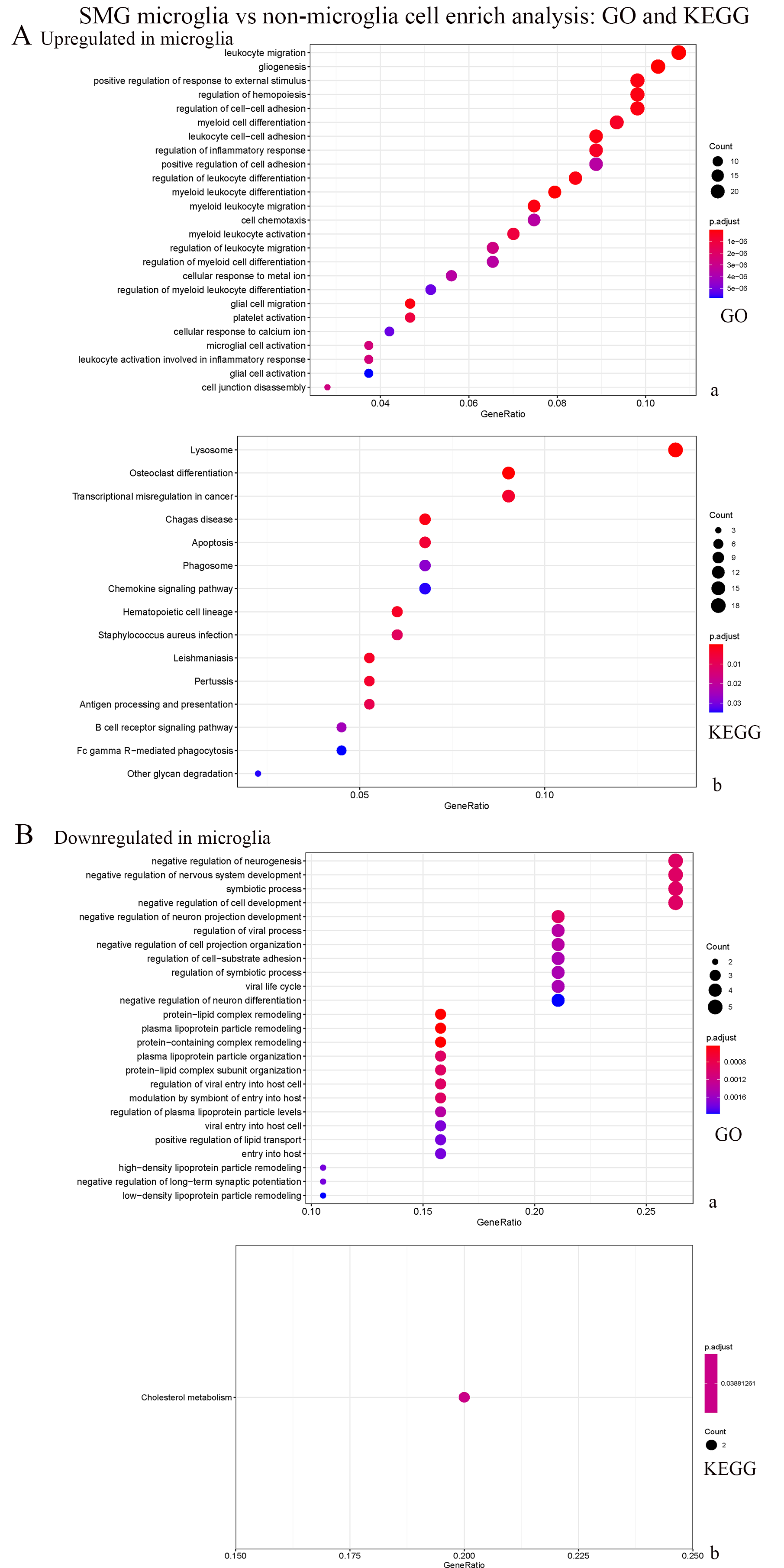

**Figure S18-** Enrichment of SMG Microglia vs non microglia cells. A: Upregulated signaling pathway in SMG; B: Downregulated signaling pathway in SMG.

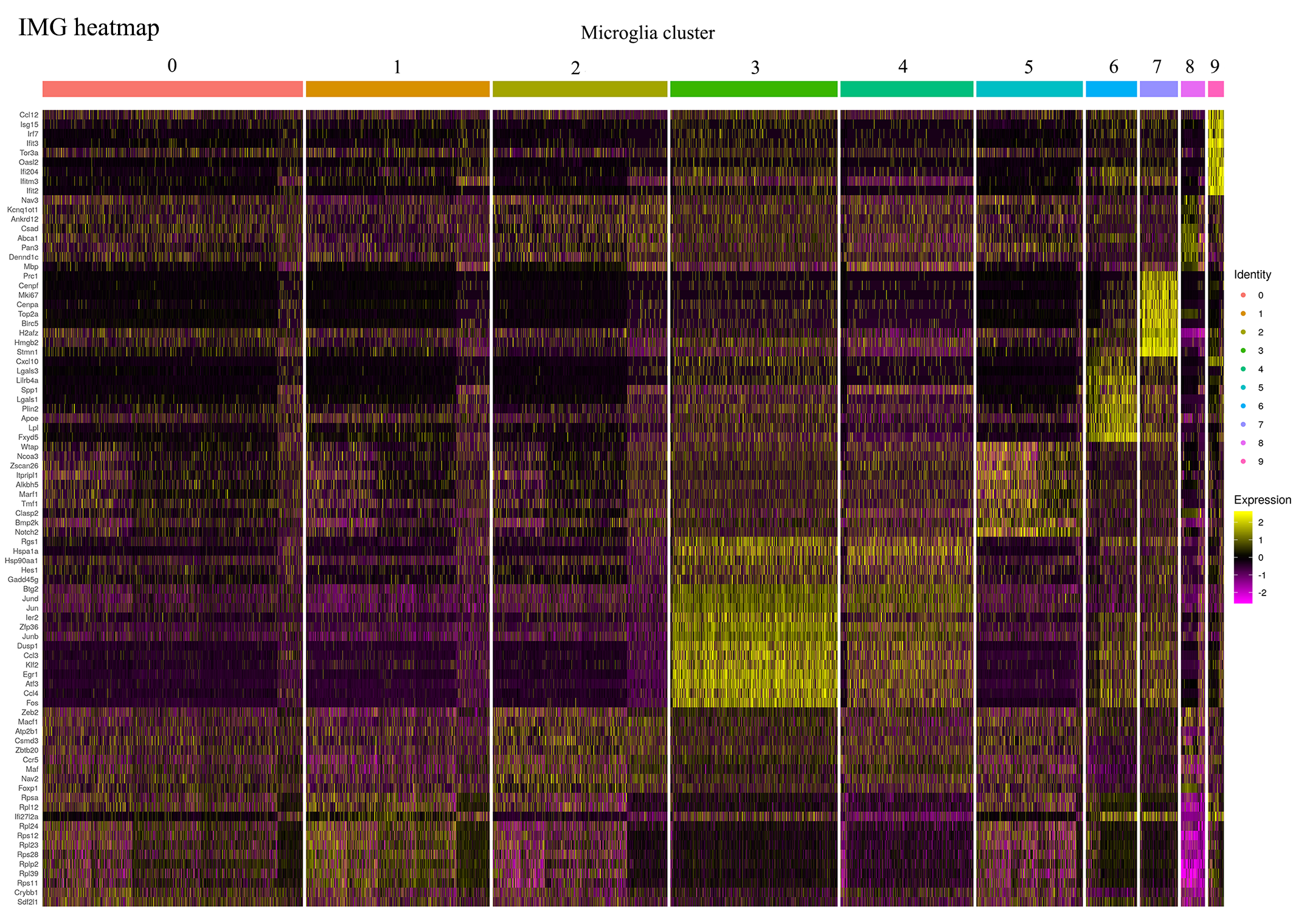

**Figure S19-** Heatmap of microglial gene expression in integration analysis. (SAH and normal microglia-IMG).

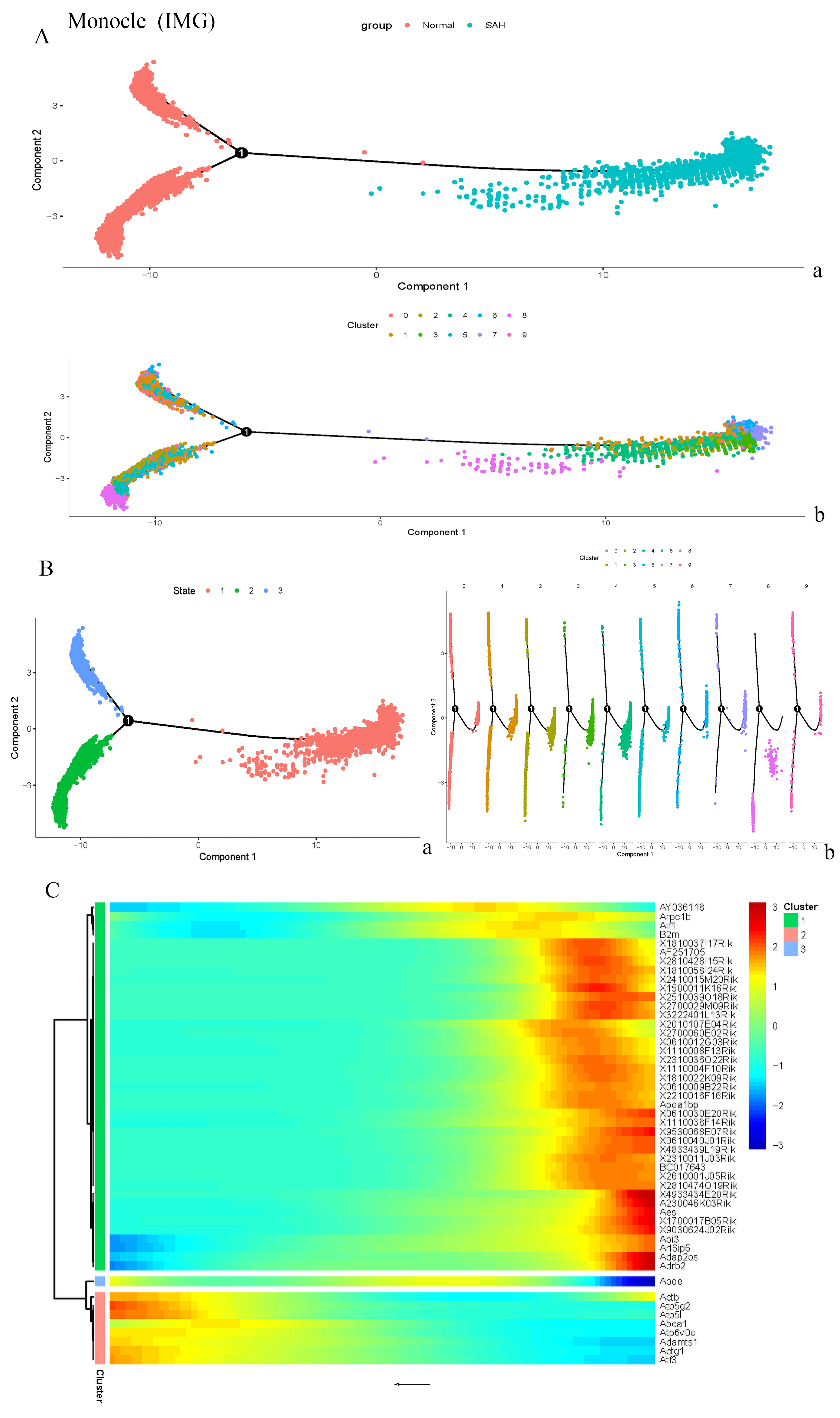

**Figure S20-** Monocle analysis of IMG. A: Monocle- state and cluster branch; B: Monocle- state branch(a), monocle cluster- split branch (b) C: Top 50 expressed genes heatmap in monocle.
